# Membrane fusion and immune evasion by the spike protein of SARS-CoV-2 Delta variant

**DOI:** 10.1101/2021.08.17.456689

**Authors:** Jun Zhang, Tianshu Xiao, Yongfei Cai, Christy L. Lavine, Hanqin Peng, Haisun Zhu, Krishna Anand, Pei Tong, Avneesh Gautam, Megan L. Mayer, Richard M. Walsh, Sophia Rits-Volloch, Duane R. Wesemann, Wei Yang, Michael S. Seaman, Jianming Lu, Bing Chen

## Abstract

The Delta variant of severe acute respiratory syndrome coronavirus 2 (SARS-CoV-2) has outcompeted previously prevalent variants and become a dominant strain worldwide. We report here structure, function and antigenicity of its full-length spike (S) trimer in comparison with those of other variants, including Gamma, Kappa, and previously characterized Alpha and Beta. Delta S can fuse membranes more efficiently at low levels of cellular receptor ACE2 and its pseudotyped viruses infect target cells substantially faster than all other variants tested, possibly accounting for its heightened transmissibility. Mutations of each variant rearrange the antigenic surface of the N-terminal domain of the S protein in a unique way, but only cause local changes in the receptor-binding domain, consistent with greater resistance particular to neutralizing antibodies. These results advance our molecular understanding of distinct properties of these viruses and may guide intervention strategies.

## Introduction

Severe acute respiratory syndrome coronavirus 2 (SARS-CoV-2) is the causative agent of the COVID-19 pandemic (*1*). Its Delta variant (*2*), first detected in India and also known as lineage B.1.617.2, is a variant of concern (VOC) and has outcompeted other previously prevalent variants to become a globally dominant strain within a few months. It has been estimated that this variant is about twice as transmissible as the virus from the initial outbreak (i.e., the strain Wuhan-Hu-1; ref(*1, 3, 4*)). Preliminary data suggest infection by the Delta variant has a shorter incubation period with a viral load of >1,000 times greater than that by earlier variants (*5*), but it remains uncertain whether or not it can cause more severe disease (*6, 7*). Nevertheless, in some individuals, the Delta variant appears to overcome immunity elicited by the first-generation vaccines designed using the Wuhan-Hu-1 sequence (*8–11*). Another VOC, Gamma (lineage B.1.1.28 or P.1), has been widespread in Brazil and some other countries (*12, 13*). A third variant, Kappa (lineage B.1.617.1), also first reported in India, remains a variant of interest (VOI) with only a limited surge in certain areas (*14, 15*). It is critical to understand the molecular mechanisms of the increased transmissibility and immune evasion of these variants to facilitate development of intervention strategies.

SARS-CoV-2 is an enveloped positive-stranded RNA virus that enters a host cell by fusion of its lipid bilayer with a membrane of the target cell. The membrane fusion reaction is catalyzed by the virus-encoded trimeric spike (S) protein after binding to the viral receptor angiotensin converting enzyme 2 (ACE2). The S protein is first produced as a single-chain precursor, which is then processed by a host furin-like protease into the receptor-binding fragment S1 and the fusion fragment S2 (Fig. S1; ref(*16*)). After engaging with ACE2 on the host cell surface, the S protein is cleaved by a second cellular protease TMPRSS2 or by cathepsins B and L (CatB/L) in S2 (S2’ site; Fig. S1; ref(*17*)), initiating dissociation of S1 and a cascade of refolding events in S2 to drive fusion of the two membranes (*18, 19*). S1 contains four domains - NTD (N-terminal domain), RBD (receptor-binding domain), and two CTDs (C-terminal domains), protecting the central helical-bundle structure formed by the prefusion S2. The RBD can adopt either a “down” conformation for a receptor-inaccessible state, or an “up” conformation for a receptor-accessible state (*20*). The RBD movement appears to be an important mechanism for the virus to protect its functionally critical receptor binding site from host immune responses (*20, 21*).

Intensive studies on the S protein have advanced our knowledge of the SARS-CoV-2 entry process substantially (reviewed in ref(*22–25*)). In the work reported here, we have characterized the full-length S proteins derived from the Delta, Kappa and Gamma variants and determined their structures by cryogenic electron microscopy (cryo-EM). Comparison of the structure, function and antigenicity of the Delta S with those of Gamma and Kappa, as well as previously characterized Alpha and Beta (*26*), provides molecular insights into mechanisms of the heightened transmissibility and enhanced immune evasion of the most contagious form of SARS-CoV-2 since its initial outbreak.

## Results

### Membrane fusion by Delta S is substantially faster than that of other variants

To characterize the full-length S proteins with the sequences derived from natural isolates of Gamma (hCoV-19/Brazil/AM-992/2020), Kappa (hCoV-19/India/MH-NEERI-NGP-40449/2021) and Delta (hCoV-19/India/GJ-GBRC619/2021) variants (Fig. S1), we transfected HEK293 cells with the respective expression constructs and compared their membrane fusion activities with that of the full-length S construct of their parental strain (G614 or B.1 variant; ref(*27*)). All S proteins expressed at comparable levels (Fig. S2A). Kappa S showed a substantially lower level of cleavage between S1 and S2 than other variants, suggesting that a P681R mutation near the furin cleavage site does not increase the S protein processing. The same mutation, also present in the Delta variant, has not significantly altered the extent of cleavage from that seen in its parent strains (Fig. S2A). The cells producing these S proteins fused efficiently with ACE2-expressing cells, as expected (Fig. S2B). The fusion activity of Kappa S was ∼50% that of other S proteins at a low transfection level, presumably due to the minimal extent of cleavage at the furin site, but the difference diminished at high transfection levels (Fig. S2B).

To test whether Delta S could fuse membranes more efficiently than other variants to account for its remarkable transmissibility, we first performed a time-course experiment with a cell-cell fusion assay, with both S and ACE2 transfected at high levels (Fig. S3A). We found no significant differences in the fusion activity among all the variants tested here, including previously characterized Alpha and Beta (*26*). We did, however, observe that Delta S-expressing cells fused with the negative-control HEK293 cells more efficiently than other variants, particularly at longer time points (Figs. 1A and S3B). HEK293 cells express a minimal level of endogenous ACE2, and they have been used as a negative control when not transfected by the ACE2 expression construct in our standard, 2-hour fusion protocol (*28*). The same pattern was also reproduced when small amounts of ACE2 were introduced in HEK293 cells, but the differences diminished when the ACE2 transfection level increased (Figs. 1B and S3C). These data suggest that Delta S can enter a host cell expressing low levels of ACE2 more efficiently than can other variants.

**Figure 1.**
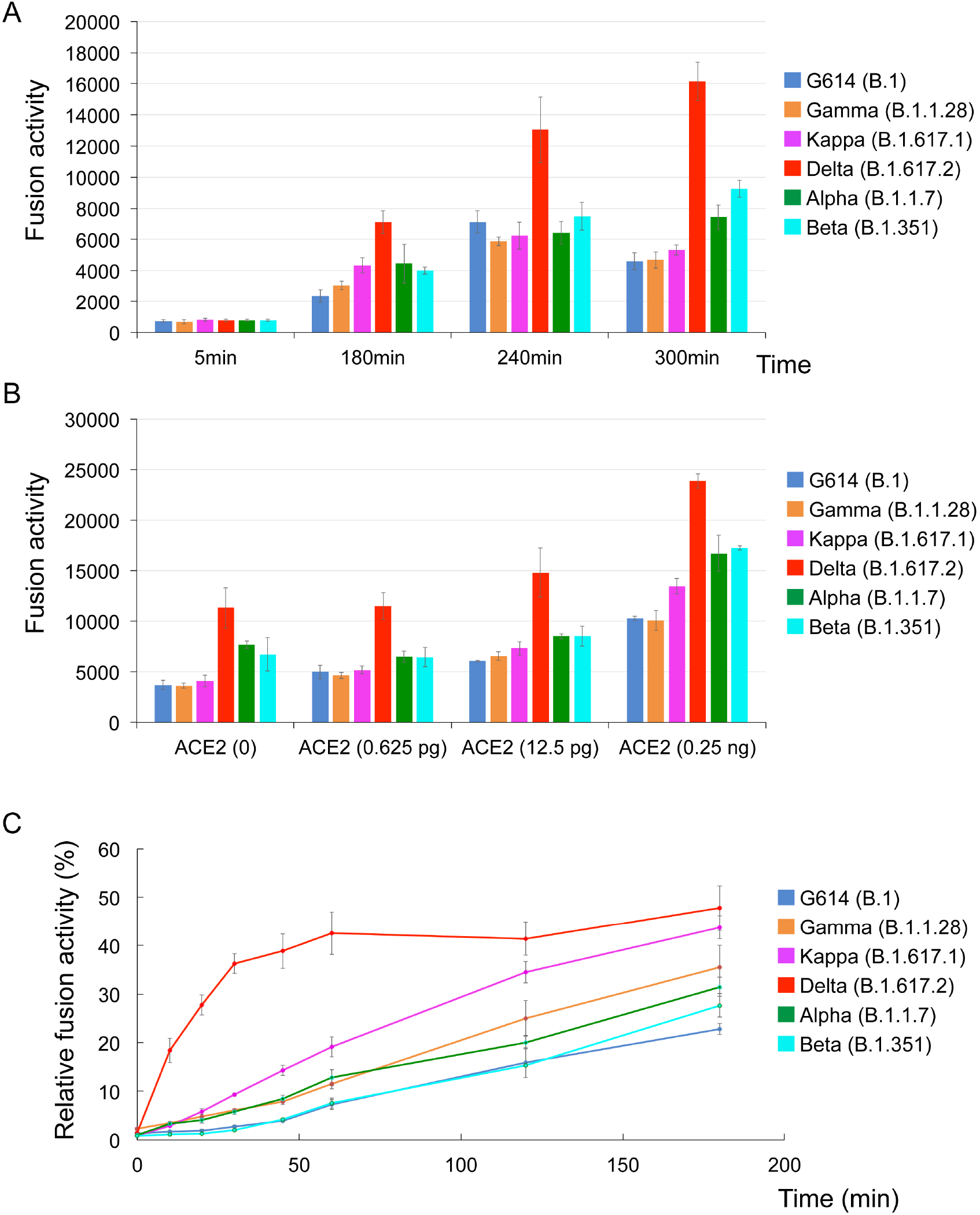
More efficient membrane fusion by the Delta variant than other variants. (**A**) Time-course of cell-cell fusion mediated by various full-length S proteins, as indicated, with HEK293 cells with no exogenous ACE2. (**B**) Cell-cell fusion mediated by various full-length S proteins with HEK293 cells transfected with low levels (0-0.25 ng) of ACE2 expression constructs. (**C**) Time course of infection HEK293-ACE2 cells by MLV-based pseudotyped viruses using various SARS-CoV-2 variant S constructs containing a CT deletion in a single cycle. Infection was initiated by mixing viruses and target cells, and viruses were washed out at each time point as indicated. The full time-course and concentration series are shown in Fig. S3. The experiments were repeated at least three times with independent samples giving similar results.

We then performed a similar time-course experiment using murine leukemia virus (MLV)-based pseudoviruses expressing the S constructs with the cytoplasmic tail deleted to facilitate incorporation into particles (*29, 30*). The infection was initiated by mixing the viruses and target cells and the viruses were washed out at each time point. The data in Figs. 1C and S3D show that the Delta variant established infection much more rapidly in the first 60-min period than did any other variant, when infectivity was normalized to its maximum level. The other variants gradually caught up over time, however, and eventually reached their maximum levels at 8 hours (Fig. S3D). Some viruses, including Delta, reproducibly showed lower measurements for the no wash-out controls than those measured at the 8-hour time point, consistent with some cytotoxicity that can reduce the expression of the reporter gene.

Our findings from both cell-based and pseudovirus-based assays suggest that the Delta variant can infect a target cell substantially more rapidly than the other variants we tested, either by more effective attachment or faster fusion kinetics.

### Biochemical and antigenic properties of the intact S proteins from the variants

To produce the full-length S proteins, we added a C-terminal strep-tag to the S proteins of the Gamma, Kappa and Delta variants (Fig. S4A), and expressed and purified these proteins by the procedures established for the D614, G614, Alpha and Beta S trimers (*26, 28, 31*). The Gamma S protein eluted in three distinct peaks, corresponding to the prefusion S trimer, postfusion S2 trimer and dissociated S1 monomer, respectively (*28*), as shown by Coomassie-stained SDS-PAGE analysis (Fig. S4B). The prefusion trimer accounted for <40% of the total protein, like the profile of the original Wuhan-Hu-1 S protein, indicating that this trimer is not very stable. Although the Kappa protein eluted in one major peak corresponding to the prefusion trimer, there is a significant amount of aggregate on the leading side and a large shoulder on the trailing side, suggesting that the protein is also not very stable and conformationally heterogenous. Moreover, a large fraction of the protein remains uncleaved (Fig. S4B), further confirming that the furin cleavage is inefficient despite the P681R mutation. In contrast, the Delta S protein eluted in a single symmetrical peak of the prefusion trimer showing little aggregation or dissociation, and it is probably the most stable S trimer preparation among all the full-length S proteins that we have examined (Fig. S3B; ref(*31*)). Negative stain EM also confirmed these results (Fig. S5). SDS-PAGE analysis showed that the Delta prefusion trimer peaks contained primarily the cleaved S1/S2 complex with a cleavage level very similar to that of the G614 and Beta S proteins (*26, 31*), again indicating little impact of the P681R mutation on the extent of furin cleavage.

To analyze antigenic properties of the prefusion S trimers, we measured their binding to soluble ACE2 proteins and S-directed monoclonal antibodies isolated from COVID-19 convalescent individuals by bio-layer interferometry (BLI). The selected antibodies recognize distinct epitopic regions on the S trimer, as defined by competing groups designated RBD-1, RBD-2, RBD-3, NTD-1, NTD-2 and S2 (Fig. S6A; ref(*32*)). The last two groups contain primarily non-neutralizing antibodies. The Gamma variant bound substantially more strongly to the receptor than did its G614 parent, regardless of the ACE2 oligomeric state (Figs. 2 and S6B; Table S1), probably because of its mutations (K417T, E484K and N501Y) in the RBD. ACE2 affinities for the Kappa and Delta S proteins were intermediate between those of the G614 and Gamma trimers, with Kappa closer to the Gamma and Delta closer to G614, except for binding of Delta S trimer with dimeric ACE2, which had an unexpectedly higher off-rate than did the other variants (Fig. 2). These data were largely confirmed by using monomeric RBD preparations instead of the S trimers of these variants, except that the Kappa RBD showed slightly higher affinity for ACE2 than the Gamma RBD (Fig. S6B and Table S1). ACE2 did not dissociate more rapidly from the Delta RBD than it did from the Gamma and Kappa RBDs; a possible explanation for the apparently weaker affinity of the Delta trimer for the ACE2 dimer may be that ACE2 binding induces S1 dissociation. Overall, these results suggest that the mutations in the RBD of the Gamma variant enhance receptor recognition, while the RBD mutations in the Kappa (L452R and E484Q) and Delta (L452R and T478K) variants have a less impact on ACE2 affinity. The dimeric ACE2 appears to be more effective in inducing S1 dissociation from the Delta S trimer than from any other variant, including Alpha and Beta (*26*).

**Figure 2.**
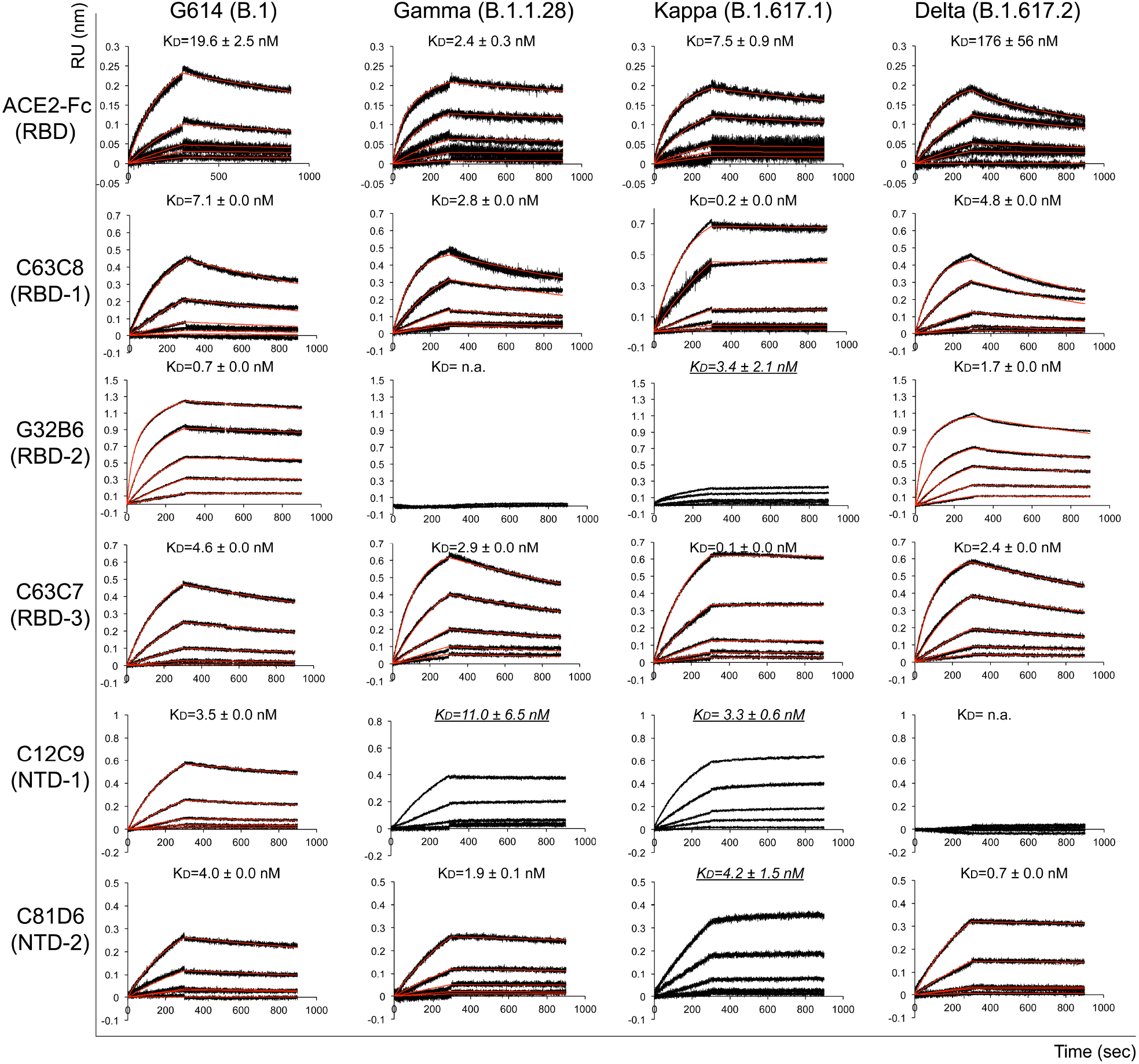
Antigenic properties of the purified full-length SARS-CoV-2 S proteins. Bio-layer interferometry (BLI) analysis of the association of prefusion S trimers derived from the G614 “parent” strain (B.1) and the Gamma (B.1.1.28), Kappa (B.1.617.1) and Delta (B.1.617.2) variants with soluble ACE2 constructs and with a panel of antibodies representing five epitopic regions on the RBD and NTD (see Fig. S4A and ref(*32*)). For ACE2 binding, purified S proteins were immobilized to AR2G biosensors and dipped into the wells containing ACE2 at various concentrations. For antibody binding, various antibodies were immobilized to AHC biosensors and dipped into the wells containing each purified S protein at different concentrations. Binding kinetics were evaluated using a 1:1 Langmuir model except for dimeric ACE2 and antibody G32B6 targeting the RBD-2, which were analyzed by a bivalent binding model. The sensorgrams are in black and the fits in red. Binding constants highlighted by underlines were estimated by steady-state analysis as described in the Methods. RU, response unit. Binding constants are also summarized here and in Table S1. All experiments were repeated at least twice with essentially identical results.

All selected monoclonal antibodies had reasonable affinities for the G614 S trimer (Figs. 2 and S6B; Table S1). The Gamma variant completely lost binding to the two RBD-2 antibodies, G32B6 and C12A2, as well as to one NTD-1 antibody, C83B6, but it could still bind another NTD-1 antibody C12C9 with somewhat reduced affinity, suggesting that these two antibodies target two overlapping but distinct epitopes. Its affinities for the remaining antibodies were the same as those of the G614 trimer. Binding of the Kappa S trimer showed unrealistically slow off-rates for a number of antibodies (Figs. 2 and S6B), presumably due to aggregation and conformational heterogeneity. Qualitatively, it had substantially weakened binding to the RBD-2 antibodies and the NTD-1 antibody C83B6, but with wildtype or even enhanced affinity for another NTD-1 antibody C12C9. Thus, the changes of its antigenic profile are similar to those of the Gamma S, but somewhat less complete. The Delta S only lost binding to the two NTD-1 antibodies with little changes in affinities for the other antibodies, including those targeting the RBD (Figs. 2 and S6B; Table S1). The BLI data were also largely consistent with the binding results with the membrane-bound S trimers measured by flow cytometry (Fig. S7).

We next analyzed the neutralization potency of these antibodies and of trimeric ACE2 (*33*) by measuring the extent to which they blocked infection by these variants in an HIV-based pseudovirus assay. For most antibodies, the neutralization potency correlated with their binding affinity for the membrane-bound or purified S proteins (Table S2). C81D6 and C163E6 recognize two non-neutralizing epitopes in the NTD and S2, respectively, and they did not neutralize any of the pseudoviruses. Thus, the mutations in the Gamma and Kappa variants have a greater impact on the antibody sensitivity of the virus than those in the Delta variant.

### Overall structures of the intact S trimers of the Delta, Kappa and Gamma variants

We determined the cryo-EM structures of the full-length S trimers with the unmodified sequences of the Delta, Kappa and Gamma variants. Cryo-EM images were acquired on a Titan Krios electron microscope equipped with a Gatan K3 direct electron detector. We used crYOLO (*34*) for particle picking, and RELION (*35*) two-dimensional (2D) classification, three dimensional (3D) classification and refinement (Figs. S8-S13). 3D classification gave three distinct classes each for both the Delta and Kappa S trimers, representing one closed prefusion conformation and two one-RBD-up conformations, respectively. There were two different classes for the Gamma trimer, representing two one-RBD-up conformations. These structures were refined to 3.1-4.4Å resolution (Fig. S8-S14; Table S3).

There are no major changes in the overall architectures of the full-length variant S proteins when compared to that of the parental G614 S trimer in the corresponding conformation (Figs. 3 and S15; ref(*31*)). In the closed prefusion conformation, the NTD, RBD, CTD1 and CTD2 of S1 wrap around the S2 trimer. In the one-RBD-up conformation, the RBD movement did not cause any changes in the central helical core structure of S2, but opened up the trimer by shifting two adjacent NTDs away from the three-fold axis of the trimer. The furin cleavage site at the S1/S2 boundary (residues 682-685), including the P681R substitution, was still not visible in any these maps.

**Figure 3.**
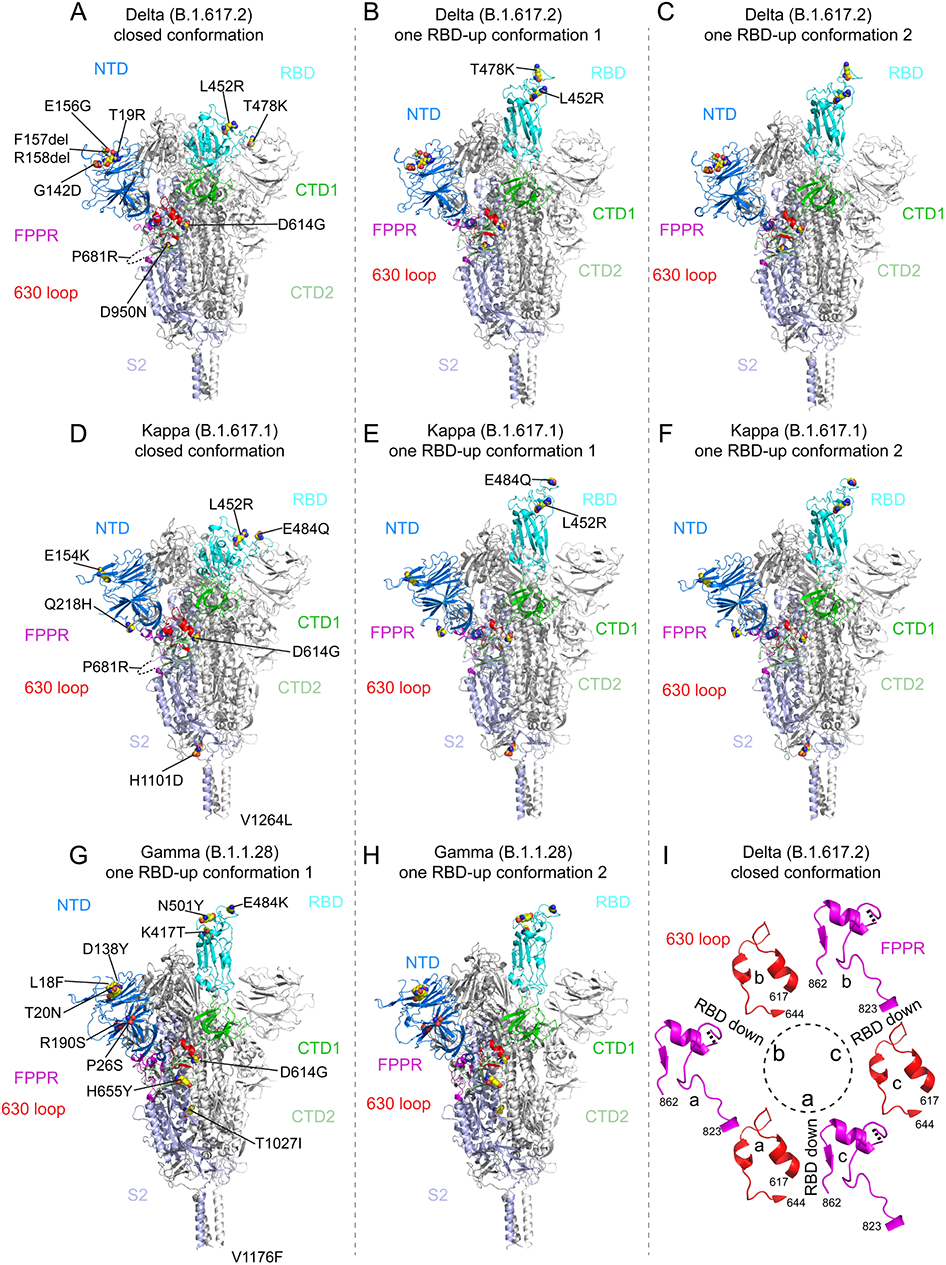
Cryo-EM structures of the full-length SARS-CoV-2 S proteins from the Delta, Kappa and Gamma variants. (**A-C**) The structures of the closed prefusion conformation and two one-RBD-up conformations of the Delta S trimer are shown in ribbon diagram with one protomer colored as NTD in blue, RBD in cyan, CTD1 in green, CTD2 in light green, S2 in light blue, the 630 loop in red and the FPPR in magenta. (**D-F**) The structures of the closed prefusion conformation and two one-RBD-up conformation of the Kappa S trimer are shown in ribbon diagram with the same color scheme as in (A). (**G**) and (**H**) The structures of the two one-RBD-up conformations of the Gamma S trimer are shown in ribbon diagram with the same color scheme as in (A). All mutations in the three variants, as compared to the original virus (D614), are highlighted in sphere model. (**I**) Structures, in the Delta closed trimer, of segments (residues 617-644) containing the 630 loop (red) and segments (residues 823-862) containing the FPPR (magenta) from each of the three protomers (a, b and c). The position of each RBD is indicated. Dashed lines indicate gaps in the chain trace (disordered loops).

We have proposed that the FPPR (fusion peptide proximal region; residues 828 to 853) and 630 loop (residues 620 to 640) are control elements and that shifts in their positions modulate the stability of the S protein and the kinetics of its structural rearrangements (*28, 31*). For the Delta and Kappa variants, the configurations of the FPPR and 630 loop are largely consistent with the distribution observed in the G614 trimer: all are structured in the RBD-down conformation, while only one the FPPR and 630-loop pair is ordered in the one-RBD-up conformations. The density of residues 841-847 in the FPPR of the Delta S in the closed prefusion state is weak probably because slight (1-2Å) downward shifts of the CTD1 and RBD, which may weaken the packing of the FPPR (Fig. S15). No class representing the closed conformation has been identified for the Gamma S from three independent data sets (Fig. S12), suggesting this conformational state is not well occupied by that variant, but one FPPR and 630-loop pair is structured in the one-RBD-up conformations of Gamma S, probably stabilizing the cleaved S trimers before receptor engagement. In all three variants, the distinct one-RBD-up structures differ only by the degree to which the up RBD and the adjacent NTD of its neighboring protomer shift away from the central threefold axis (Fig. S12). Density for an N-linked glycan at residue Asn343 in the RBD has become stronger in maps of all the new variants, particularly Delta and Kappa, than in that of the G614 trimer (Fig. S16). The distal end of the glycan appears to contact the neighboring RBD, forming a ring-like density to help clamp down the three RBDs. Nonetheless, it remains unclear why the Gamma prefusion trimer dissociates, the Kappa trimer tends to aggregate and the Delta trimer is the most stable of the three.

### Structural consequences of mutations in the Delta variant

We superposed the structures of the Delta S trimer onto the G614 trimer in the closed conformation aligning them by the S2 region (Fig. S12), revealing the most prominent differences in the NTD, which contains three point mutations (T19R, G142D and E156G) and a two-residue deletion (F157del and R158del). When the two NTDs are aligned (Fig. 4A and 4B), it appears that the mutations reshape the 143-154 loop, projecting it away from the viral membrane together with an N-linked glycan (N149). They also reconfigure the N-terminal segment and the 173-187 loop, substantially altering the antigenic surface near the NTD-1 epitope group in the NTD. These structural changes are fully consistent with loss of binding and neutralization by NTD-1 antibodies (Figs. 2 and S6; Table S2). There are two mutations, L452R and T478K, in the Delta RBD, which do not lead to any major structural rearrangements in the domain (Fig. 4C). These two residues are not in the ACE2 contacting surface, and it is therefore not surprising they have little impact on the receptor binding (Fig. S17). Neither binding nor neutralization of the Delta variant by most anti-RBD antibodies tested here have changed, suggesting that the two residues are not in any major neutralizing epitopes either.

**Figure 4.**
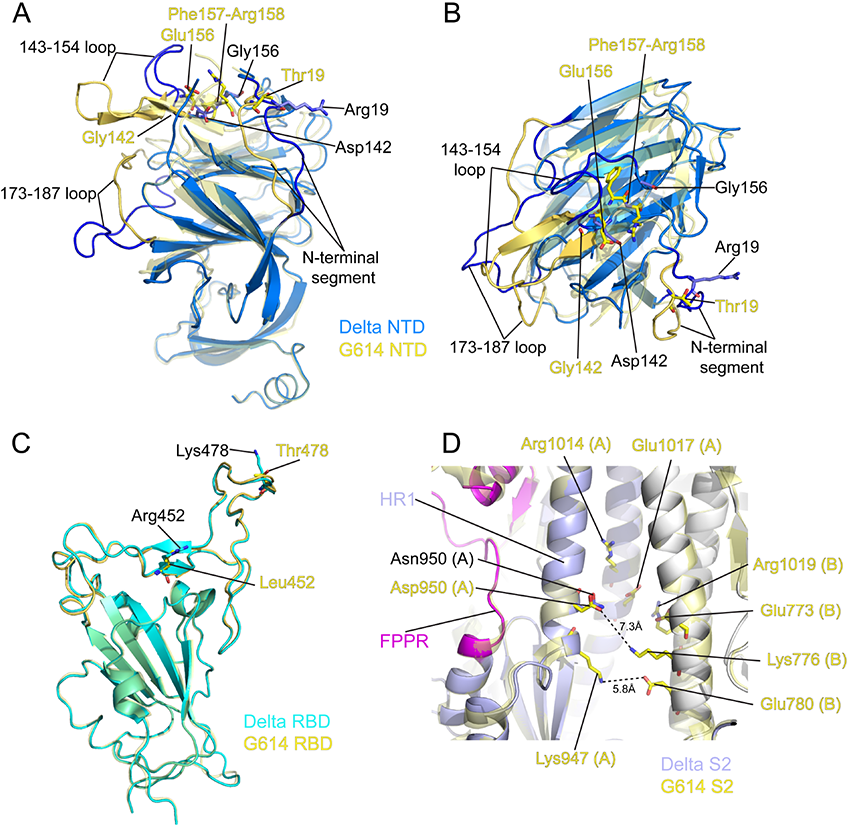
Structural impact of the mutations in the Delta S. (**A**) Superposition of the NTD structure of the Delta S trimer in blue with the NTD of the G614 S trimer (PDB ID: 7KRQ) in yellow. Locations of mutations T19R, G142D, E156G and deletion of F157 and R158 are indicated and these residues are shown in stick model. The N-terminal segment, 143-154 and 173-187 loops are rearranged between the two structures and highlighted in darker colors. (**B**) Top view of the panel (A). (**C**) Superposition of the RBD structure of the Delta S trimer in cyan with the RBD of the G614 S trimer in yellow. Locations of mutations L452R and T478K are indicated and these residues are shown in stick model. (**D**) A close-up view of superposition of the Delta S2 in light blue with the S2 of the G614 S trimer in yellow near residue 950. Locations of the D950N mutation and charged residues in the vicinity including Lys947, Arg1014, Glu1017 from the protomer A and Glu773, Lys776, Glu780 and Arg1019 from the protomer B are indicated. All these residues are shown in stick model.

We can detect no obvious structural alterations from the substitution D950N in S2 (Fig. 4D). This residue is in HR1 (heptad repeat 1) and also close to the FPPR. Although D950 in the G614 trimer is not close enough to form a salt bridge with any positively charged residue nearby, there are multiple pairs of charged residues in the vicinity that could help stabilize the packing between S2 protomers in the prefusion conformation. Thus, modifying the local electrostatic potential by the D950N substitution might introduce subtle changes that can influence the conformational changes of S2 required for membrane fusion.

### Structural impact of the mutations in the Kappa and Gamma variants

There are only two mutations (E154K and Q218H) in the NTD of the Kappa variant (Fig. 5A). In the G614 trimer, Glu154 forms a salt bridge with Arg102 (*31*). E154K substitution not only eliminates the ionic interaction, but also exerts a repulsive force on Arg102, possibly impacting the nearby 173-187 loop, which is disordered in the Kappa trimer. Residue 218 is surface-exposed and on the opposite side from the neutralizing epitopes. Q218H appeared to shift the adjacent 210-217 loop and perhaps also contributed to the rearrangement of the 173-187 loop (Fig. 5A). Like the Delta RBD, there are two mutations (L452R and E484Q) in the Kappa RBD (Fig. 5B), which do not alter the overall structure of the domain. Glu484 can form a salt bridge with ACE2 Lys31 in the RBD-ACE2 complex (Fig. S17; ref(*36, 37*)). The E484Q substitution loses the salt bridge, but hydrogen bonds between Glu484 and ACE2 Lys31 might compensate and thus account for a small increase in ACE2 binding affinity. L452R, also present in the Delta RBD, does not seem to have a significant impact on ACE2 binding (Fig. S17). In addition, the mutation H1101D in S2 caused little local changes (Fig. S18A), and V1264L is not visible in our structures.

**Figure 5.**
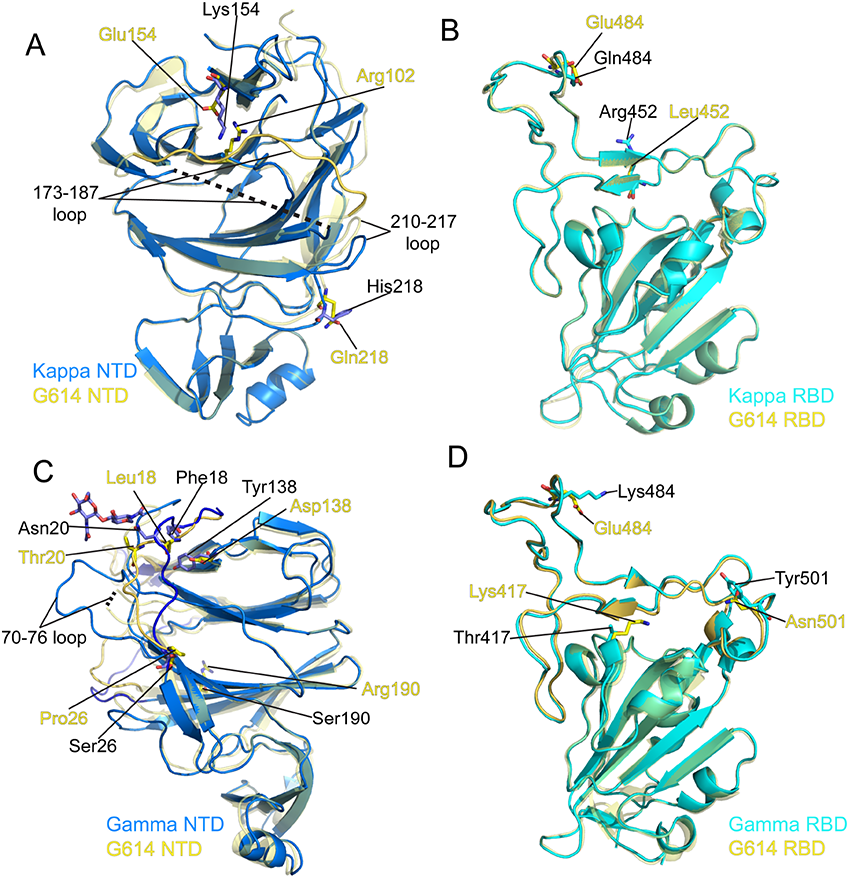
Structural impact of the mutations in the Kappa and Gamma S proteins. (**A**) Superposition of the NTD structure of the Kappa S trimer in blue with the NTD of the G614 S trimer in yellow. Locations of mutations E154K and Q218H, as well as Arg102 that forms a salt bridge with Glu154 in the G614 structure are indicated and these residues are shown in stick model. The 173-187 loop in the G614 trimer is highlighted in a darker color; it becomes disordered in the Kappa trimer. (**B**) Superposition of the RBD structure of the Kappa S trimer in cyan with the RBD of the G614 S trimer in yellow. Locations of mutations L452R and E484Q are indicated and these residues are shown in stick model. (**C**) A view of superposition of the NTD structures of the Gamma (blue) and G614 (yellow; PDB ID: 7KRR) S trimers in the one-RBD-up conformation. Locations of mutations L18F, T20N, P26S, D138Y and R190S, as well as N-linked glycan attached to Asn20 in the Gamma structure are indicated and these residues are shown in stick model. (**D**) Superposition of the RBD structure of the Gamma S trimer in cyan with the RBD of the G614 S trimer in yellow. Locations of mutations K417T, E484K and N501Y are indicated and these residues are shown in stick model.

We collected three large independent data sets for the Gamma S protein, but the number of good particles that could be extracted remained relatively low because of the instability and heterogeneity of the protein sample, giving two maps at 3.8Å and 4.4Å resolution, respectively (Figs. S12 and S13). Nonetheless, the structural changes in the NTD caused by the mutations (L18F, T20N, P26S, D138Y and R190S) were evident even in these maps. All mutations except for R190S are near the N-terminal segment and contribute to reconfigure its extended structure (Fig. 5C). The new conformation of the N-terminal segment appears to stabilize the 70-76 loop, disordered in most of SARS-CoV-2 S trimer structures published previously (*20, 28, 38*). T20N has created a new glycosylation site and Asn20 is indeed glycosylated in the Gamma NTD protein (Fig. 5C). These changes apparently also shift the 143-154 and 173-187 loops in the Gamma variant (Fig. S18B), leading to a relatively large-scale rearrangement of the antigenic surface of the NTD. Like the Beta RBD (*26, 39*), Gamma RBD has three mutations, K417T, E484K and N501Y, which do not produce any major structural rearrangements (Fig. 5D). N501Y increases receptor-binding affinity, which may be counteracted by K417T and E484K because of loss of ionic interactions with ACE2 (Fig. S17; ref(*26, 39*)). K417T and E484K are probably responsible for loss of binding and neutralization of the Gamma by antibodies that target the RBD-2 epitopes. H655Y in the CTD2 did not change the local structure (Fig. S18C), but its location near the N-terminus of the cleaved S2 suggest that this mutation may play a role in destabilizing the S trimer of this variant. Finally, T1027I did not lead to any major changes in S2 (Fig. S18D), and V1176F is in a disordered region.

## Discussion

The Delta variant of SARS-CoV-2 has rapidly replaced the previously dominant variants, including Alpha, which is itself ∼60% more transmissible than the original Wuhan-Hu-1 strain (*40–42*). Delta thus appears to have acquired enhanced capacity for propagating in human cells. Several hypotheses have been proposed to explain its heighten transmissibility, including mutations in the RBD enhancing receptor engagement (*43*), P681R substitution near the S1/S2 boundary leading to more efficient furin cleavage (*44, 45*), and changes in its RNA polymerase increasing viral replication. We can not rule out the possibility that mutations in the viral replication machinery unique to the Delta variant (e.g. G671S in nsp12) may greatly increase the production of genomic RNA, but viral assembly into mature virions would require many other factors to achieve the >1,000 fold greater viral load in infected patients. We have not detected any significant increase in ACE2 binding by either the full-length Delta S trimer or its RBD fragment, nor have we observed more efficient cleavage in the Delta S than any other variants. Indeed, the P681R mutation is also present in the Kappa variant, which appears to have impaired furin cleavage, at least, in HEK293 cells used in our experiments. We have identified two properties, apparently unique to the Delta variant among those that have been studied so far, that might possibly account for its unusual transmissibility. First, when the Delta S protein is expressed on the cell surface at a saturating level, those cells fuse more efficiently with target cells that produce low levels of ACE2 than do cells of any other variant, including previously characterized Alpha and Beta (*26*). When the ACE2 expression level increases, the differences among the variants diminish. Second, the pseudoviruses containing the Delta S construct enter the ACE2-expressing target cells substantially more rapidly than other variants. These data suggest that the Delta S protein has evolved to optimize the fusion step for entering cells expressing low levels of the receptor. This optimization may explain why the Delta variant can transmit upon relatively brief exposure and infect many more host cells rapidly, leading to a short incubation period and greater viral load during the infection. A caveat is that all our experiments were performed in vitro; additional studies with authentic viruses will be needed to confirm our findings in more clinically relevant settings.

If our hypothesis is valid, what is the structural basis for the enhanced fusogenicity of the Delta S protein? All mutations but one in the Delta S are located in either the RBD or NTD. Our extensive binding studies indicate that the Delta S does not engage the receptor ACE2 more tightly than does any other variant. It is unclear what other functional roles the NTD may play in the membrane fusion process, besides protecting the nearby RBDs. If the mutations in the NTD enhance RBD exposure to potential receptors, we should have observed, in our cryo-EM study, more particles in the RBD-up conformation from the Delta data set. Thus, the structural changes in both the RBD and NTD are unlikely to explain the efficient membrane fusion by the Delta variant. The last mutation, D950N, is unique to Delta and located in HR1 of S2 near the FPPR. D950N eliminates a negative charge (three in a trimer), but we have not observed any obvious structural changes caused by this substitution in the prefusion conformation. Its location nonetheless appears to be a critical site that can influence the refolding of S2, required for membrane fusion. Although D950 is not involved in a salt bridge in the G614 trimer, it is conceivable that the local change in the electrostatic potential may destabilize the prefusion S2 in a very subtle way because there are several pairs of charged residues in the vicinity. Indeed, too much destabilization of the prefusion conformation may be detrimental since it may prompt the S trimer to undergo premature conformational changes and inactivate the protein. Thus, successful viral evolution must be a delicate balancing act to avoid tampering with its role in fusion. Future studies are clearly needed to fully address the question.

The RBD and NTD are the two major sites on the S trimer targeted by neutralizing antibodies characterized previously (*32, 46–48*). The three strains studied here show once again how different variants can use different strategies to remodel their NTD and evade host immunity. One important implication is that the function of the NTD does not require specific structural elements or sequences since the surface loops, β strands in the core structure and even some N-linked glycans can be rearranged in different ways without compromising viral infectivity. In contrast, the overall structure of the RBD has been strictly preserved among all the variants and reoccurring surface mutations appear to be limited to a number of sites (e.g. K417 in AY1 and AY2 sublineages of the Delta variant (*49*)), consistent with its critical role in receptor binding. We therefore suggest that therapeutic antibodies or universal vaccines should not target the NTD since escaping from anti-NTD antibodies appears to be at little cost to the virus.

Continuous spread of SARS-CoV-2 worldwide will inevitably allow emergence of new variants as the virus evolves to survive under the selective pressure exerted by increasingly prevalent host immunity at the population level. Indeed, new sublineages of the Delta variant have already been detected, including AY.1, AY.2 and AY.3 (*49*). How to address the viral diversity will be the biggest challenge for development of next-generation vaccines and therapeutics. Our structure, function and antigenicity studies of the SARS-CoV-2 Delta, Kappa and Gamma variants show the molecular events that have led these viruses to adapt in human communities and to evade host immunity. Our analysis suggests a mechanism for the heightened transmissibility of the most contagious variant since the beginning of the SARS-CoV-2 outbreak. The results may guide future intervention strategies.

## Materials and Methods

### Expression constructs

Genes of full-length spike (S) protein from Gamma (hCoV-19/Brazil/AM-992/2020; GISAID accession ID: EPI_ISL_833172), Kappa (hCoV-19/India/MH-NEERI-NGP-40449/2021; GISAID accession ID: EPI_ISL_1547802) and Delta (hCoV-19/India/GJ-GBRC619/2021; GISAID accession ID: EPI_ISL_2020954) were synthesized by Twist Bioscience (South San Francisco, CA) or GENEWIZ (South Plainfield, NJ). The S genes were fused with a C-terminal twin Strep tag (SGGGSAWSHPQFEKGGGSGGGSGGSSAWSHPQFEK) and cloned into a mammalian cell expression vector pCMV-IRES-puro (Codex BioSolutions, Inc, Gaithersburg, MD).

### Expression and purification of recombinant proteins

Expression and purification of the full-length S proteins were carried out as previously described (*28*). Briefly, expi293F cells (ThermoFisher Scientific, Waltham, MA) were transiently transfected with the S protein expression constructs. To purify the S protein, the transfected cells were lysed in a solution containing Buffer A (100 mM Tris-HCl, pH 8.0, 150 mM NaCl, 1 mM EDTA) and 1% (w/v) n-dodecyl-β-D-maltopyranoside (DDM) (Anatrace, Inc. Maumee, OH), EDTA-free complete protease inhibitor cocktail (Roche, Basel, Switzerland), and incubated at 4°C for one hour. After a clarifying spin, the supernatant was loaded on a strep-tactin column equilibrated with the lysis buffer. The column was then washed with 50 column volumes of Buffer A and 0.3% DDM, followed by additional washes with 50 column volumes of Buffer A and 0.1% DDM, and with 50 column volumes of Buffer A and 0.02% DDM. The S protein was eluted by Buffer A containing 0.02% DDM and 5 mM d-Desthiobiotin. The protein was further purified by gel filtration chromatography on a Superose 6 10/300 column (GE Healthcare, Chicago, IL) in a buffer containing 25 mM Tris-HCl, pH 7.5, 150 mM NaCl, 0.02% DDM. All RBD proteins were purchased from Sino Biological US Inc (Wayne, PA).

The monomeric ACE2 or dimeric ACE2 proteins were produced as described (*33*). Briefly, Expi293F cells transfected with monomeric ACE2 or dimeric ACE2 expression construct and the supernatant of the cell culture was collected. The monomeric ACE2 protein was purified by affinity chromatography using Ni Sepharose excel (Cytiva Life Sciences, Marlborough, MA), followed by gel filtration chromatography. The dimeric ACE2 protein was purified by GammaBind Plus Sepharose beads (GE Healthcare), followed gel filtration chromatography on a Superdex 200 Increase 10/300 GL column. All the monoclonal antibodies were produced as described (*32*).

### Western blot

Western blot was performed using an anti-SARS-COV-2 S antibody following a protocol described previously (*50*). Briefly, full-length S protein samples were prepared from cell pellets and resolved in 4-15% Mini-Protean TGX gel (Bio-Rad, Hercules, CA) and transferred onto PVDF membranes. Membranes were blocked with 5% skimmed milk in PBS for 1 hour and incubated a SARS-CoV-2 (2019-nCoV) Spike RBD Antibody (Sino Biological Inc., Beijing, China, Cat: 40592-T62) for another hour at room temperature. Alkaline phosphatase conjugated anti-Rabbit IgG (1:5000) (Sigma-Aldrich, St. Louis, MO) was used as a secondary antibody. Proteins were visualized using one-step NBT/BCIP substrates (Promega, Madison, WI).

### Negative stain EM

To prepare grids, 3 µl of freshly purified full-length S protein was adsorbed to a glow-discharged carbon-coated copper grid (Electron Microscopy Sciences), washed with deionized water, and stained with freshly prepared 1.5% uranyl formate. Images were recorded at room temperature at a magnification of 67,000x and a defocus value of 2.5 µm following low-dose procedures, using a Tecnai T12 electron microscope (Thermo Fisher Scientific) equipped with a Gatan UltraScan 895 4k CCD camera and operated at a voltage of 120 keV.

### Cell-cell fusion assay

The cell-cell fusion assay, based on the α-complementation of E. coli β-galactosidase, was used to measure fusion activity of SARS-CoV2 S proteins, as described (*28*). Briefly, HEK293T cells were transfected by polyethylenimine (PEI) (80 µg) with various amounts of the full-length SARS-CoV2 (Wuhan-Hu-1, G614, Alpha, Beta, Gamma, Delta or Kappa) S construct, as indicated in each specific experiment (0.025-10 µg), and the α fragment of E. coli β-galactosidase construct (10 µg), as well as the empty vector to make up the total DNA amount to 20 µg, to generate S-expressing cells. The full-length ACE2 construct, as indicated in each specific experiment (0.625 pg-10 µg) together with the ω fragment of E. coli β-galactosidase construct (10 µg), and the empty vector when needed were used to transfect HEK293T cells to create target cells. After incubation at 37°C for 24 hrs, the cells were detached using PBS buffer and resuspended in complete DMEM medium. 50 µl S-expressing cells (1.0×10^6^ cells/ml) were mixed with 50 µl ACE2-expressing target cells (1.0×10^6^ cells/ml) to allow cell-cell fusion to proceed at 37°C for from 5 min to 6 hours as indicated. Cell-cell fusion activity was quantified using a chemiluminescent assay system, Gal-Screen (Applied Biosystems, Foster City, CA), following the standard protocol recommended by the manufacturer. The substrate was added to the cell mixture and allowed to react for 90 min in dark at room temperature. The luminescence signal was recorded with a Synergy Neo plate reader (Biotek, Winooski, VT).

### Binding assay by bio-layer interferometry (BLI)

Binding of monomeric or dimeric ACE2 to the full-length Spike protein of each variant was measured using an Octet RED384 system (ForteBio, Fremont, CA), following the protocol described previously (*33*). Briefly, a full-length S protein was immobilized to Amine Reactive 2nd Generation (AR2G) biosensors (ForteBio, Fremont, CA) and dipped in the wells containing the ACE2 protein at various concentrations (5.56-450 nM for monomeric ACE2; 0.926-75 nM for dimeric ACE2) for association for 5 minutes, followed by a 10 min dissociation phase in a running buffer (PBS, 0.02% Tween 20, 2 mg/ml BSA). To measure binding of a full-length S protein to monoclonal antibodies, the antibody was immobilized to anti-human IgG Fc Capture (AHC) biosensor (ForteBio, Fremont, CA) following a protocol recommended by the manufacturer. The full-length S protein was diluted using a running buffer (PBS, 0.02% Tween 20, 0.02% DDM, 2 mg/ml BSA) to various concentrations (0.617-50 nM) and transferred to a 96-well plate. The sensors were dipped in the wells containing the S protein solutions for 5 min, followed with a 10 min dissociation phase in the running buffer. Control sensors with no S protein or antibody were also dipped in the ACE2 or S protein solutions and the running buffer as references. Recorded sensorgrams with background subtracted from the references were analyzed using the software Octet Data Analysis HT Version 12.0 (ForteBio). Binding kinetics was evaluated using a 1:1 Langmuir model except for dimeric ACE2 and antibodies G32B6 and C12A2, which were analyzed by a bivalent binding model. Sensorgrams showing unrealistic off-rates were fit individually to a single exponential function shown below to obtain the steady state response R_eq_ at each concentration.

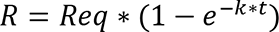

The Kd was obtained by fitting R_eq_ value and its corresponding concentration to the model: “one site-specific” using GraphPad Prism 8.0.2 according to H.J. Motulsky, Prism 5 Statistics Guide, 2007, GraphPad Software Inc., San Diego CA, www.graphpad.com).

### Flow cytometry

Expi293F cells (ThermoFisher Scientific) were grown in Expi293 expression medium (ThermoFisher Scientific). Cell surface display DNA constructs for the SARS-CoV-2 spike variants together with a plasmid expressing blue fluorescent protein (BFP) were transiently transfected into Expi293F cells using ExpiFectamine 293 reagent (ThermoFisher Scientific) per manufacturer’s instruction. Two days after transfection, the cells were stained with primary antibodies or the histagged ACE2_615_-foldon T27W protein (*33*) at 10 µg/ml concentration. For antibody staining, an Alexa Fluor 647 conjugated donkey anti-human IgG Fc F(ab’)2 fragment (Jackson ImmunoResearch, West Grove, PA) was used as secondary antibody at 5 µg/ml concentration. For ACE2_615_-foldon T27W staining, APC conjugated anti-HIS antibody (Miltenyi Biotec, Auburn, CA) was used as secondary antibody at 1:50 dilution. Cells were run through an Intellicyt iQue Screener Plus flow cytometer. Cells gated for positive BFP expression were analyzed for antibody and ACE2_615_-foldon T27W binding. The flow cytometry assays were repeated three times with essentially identical results.

### MLV-based pseudovirus assay

Murine Leukemia Virus (MLV) particles (plasmids of the MLV components kindly provided by Dr. Gary Whittaker at Cornell University and Drs. Catherine Chen and Wei Zheng at National Center for Advancing Translational Sciences, National Institutes of Health), pseudotyped with various SARS-CoV-2 S protein constructs, were generated in HEK 293T cells, following a protocol described previously for SARS-CoV (*51, 52*). To enhance incorporation of S protein into the particles, the C-terminal 19 residues in the cytoplasmic tail of each S protein were deleted. To prepare for infection, 7.5×10^3^ of HEK 293 cells, stably transfected with a full-length human ACE2 expression construct, in 15 μl culture medium were plated into a 384-well white-clear plate coated with poly-D-Lysine to enhance the cell attachment. On day 2, 15 μl of MLV pseudoviruses for each variant were added into each well pre-seeded with HEK293-ACE2 cells. The plate was centrifuged at 114 xg for 5 min at 12°C. After incubation of the pseudoviruses with the cells for a time period (10 min-8 hr), as indicated in the figures, the medium was removed and the cells were washed once with 1xDPBS. 30 μl of fresh medium was added back into each well. The cells were then incubated at 37°C for additional 40 hr. Luciferase activities were measured with Firefly Luciferase Assay Kit (CB-80552-010, Codex BioSolutions Inc).

### HIV-based pseudovirus assay

Neutralizing activity against SARS-CoV-2 pseudovirus was measured using a single-round infection assay in 293T/ACE2 target cells. Pseudotyped virus particles were produced in 293T/17 cells (ATCC) by co-transfection of plasmids encoding codon-optimized SARS-CoV-2 full-length S constructs, packaging plasmid pCMV DR8.2, and luciferase reporter plasmid pHR’ CMV-Luc. G614 S, packaging and luciferase plasmids were kindly provided by Dr. Barney Graham (Vaccine Research Center, NIH). The 293T cell line stably overexpressing the human ACE2 cell surface receptor protein was kindly provided by Drs. Michael Farzan and Huihui Ma (The Scripps Research Institute). For neutralization assays, serial dilutions of monoclonal antibodies (mAbs) were performed in duplicate followed by addition of pseudovirus. Pooled serum samples from convalescent COVID-19 patients or pre-pandemic normal healthy serum (NHS) were used as positive and negative controls, respectively. Plates were incubated for 1 hour at 37°C followed by addition of 293/ACE2 target cells (1×10^4^/well). Wells containing cells + pseudovirus (without sample) or cells alone acted as positive and negative infection controls, respectively. Assays were harvested on day 3 using Promega BrightGlo luciferase reagent and luminescence detected with a Promega GloMax luminometer. Titers are reported as the concentration of mAb that inhibited 50% or 80% virus infection (IC_50_ and IC_80_ titers, respectively). All neutralization experiments were repeated twice with similar results.

### Cryo-EM sample preparation and data collection

To prepare cryo EM grids, 3.5 µl of the freshly purified sample from the peak fraction in DDM at ∼2.5 mg/ml for the Delta variant or ∼2.0 mg/ml for the Gamma and Kappa variants was applied to a 1.2/1.3 Quantifoil grid (Quantifoil Micro Tools GmbH), which had been glow discharged with a PELCO easiGlow^TM^ Glow Discharge Cleaning system (Ted Pella, Inc.) for 60 s at 15 mA. Grids were immediately plunge-frozen in liquid ethane using a Vitrobot Mark IV (ThermoFisher Scientific), and excess protein was blotted away by using grade 595 filter paper (Ted Pella, Inc.) with a blotting time of 4 s, a blotting force of −12 at 4°C with 100% humidity. The grids were first screened for ice thickness and particle distribution. Selected grids were used to acquire images by a Titan Krios transmission electron microscope (ThermoFisher Scientific) operated at 300 keV and equipped with a BioQuantum GIF/K3 direct electron detector. Automated data collection was carried out using SerialEM version 3.8.6 (*53*) at a nominal magnification of 105,000× and the K3 detector in counting mode (calibrated pixel size, 0.825 Å) at an exposure rate of 20.24 (for Delta), ∼20.69/20.63/27.13 (for three data sets of Gamma), or ∼21.12/20.10 (for two data sets of Kappa) electrons per pixel per second. Each movie add a total accumulated electron exposure of ∼51.48 (Delta), ∼54.72/54.56/53.4 (Gamma), or ∼51.63/51.15 (Kappa) e-/Å^2^, fractionated in 50 (Delta), 51/51/50 (Gamma), or 51/51 (Kappa) frames. Data sets were acquired using a defocus range of 0.8-2.2 (Delta), 0.8-2.3 (Gamma), or 0.8-2.2 (Kappa) µm.

### Image processing and 3D reconstructions

Drift correction for cryo-EM images was performed using MotionCor2 (*54*), and contrast transfer function (CTF) was estimated by Gctf (*55*) using motion-corrected sums without dose-weighting. Motion corrected sums with dose-weighting were used for all other image processing. crYOLO was used for particle picking and RELION3.0.8 for 2D classification, 3D classification and refinement procedure. For the Delta sample, 1,830,328 particles were extracted from 20,274 images using crYOLO with a trained model, and then subjected to 2D classification, giving 1,386,630 good particles. A low-resolution negative-stain reconstruction of the Wuhan-Hu-1 (D614) sample was low-pass filtered to 40Å resolution and used as an initial model for 3D classification with C1 symmetry. After two rounds of 3D classification, two major classes with clear structural features were combined and subjected to a third round of 3D classification in C1 symmetry. The calculation led to five major classes, one in a closed, three RBD-down conformation, and four in the one-RBD-up conformation. The class of the closed conformation was re-extracted unbinned and subjected to one round of 3D auto-refinement, giving a map at 4.2Å resolution from 125,763 particles. Additional round of signal-subtraction and 3D classification using a mask for the apex region of the S trimer were performed, leading to three distinct classes, two in the closed conformation and one in the one-RBD-up conformation. The two classes in the closed conformation were combined and subjected to another round of 3D auto-refinement, followed by CTF refinement, particle polishing, and 3D auto-refinement, producing a map at 4.2Å resolution from 102,521 particles. Second round of signal-subtraction and 3D classification focusing on the apex region were carried out, giving one major class in the closed conformation with 94,680 particles. Further round of 3D auto-refinement in C3 symmetry using a whole mask was applied to this class, followed by CTF refinement, particle polishing, and 3D auto-refinement, yielding a map at 3.1Å resolution. Four classes in the one-RBD-up conformation from the third rounds of 3D classification were combined and particles re-extracted unbinned for one round of 3D auto-refinement, then further combined with the RBD-up class from the first round of signal-subtraction/ classification based on the apex region of the closed S trimer. This combined class containing 255,909 particles was autorefined producing a map at 4.1Å resolution. Particle CTF refinement/particle polishing and another round of autorefinement were performed before subjected to signal-subtraction and 3D classification using the apex mask. Two major classes in the one-RBD-up conformation were produced and they were subjected to 3D auto-refinement in C1 symmetry using a whole mask, CTF refinement, particle polishing and a final round of 3D auto-refinement. The final reconstructions have 191,067 and 25,370 particles with maps 3.4Å and 4.3Å resolution, respectively. Additional rounds of 3D auto-refinement were performed for each class using different sizes of masks at the top region to improve local resolution. The best density maps were used for model building.

For the Kappa sample, two data sets were collected and initially processed separately. 1,199,999 and 1,718,600 particles were extracted using crYOLO with a trained model from 22,019 and 17,314 images, respectively, from the two sets. The selected particles were subjected to 2D classification, giving 1,112,384 and 1,649,296 particles, respectively. A low-resolution negative-stain reconstruction of the Wuhan-Hu-1 (D614) sample was low-pass-filtered to 40Å as an initial model for 3D classification with C1 symmetry. Three rounds of 3D classification were performed separately for each data set. For data set 1, two major classes were identified, one in the closed conformation and the other in the one-RBD-up conformation. The class in the closed state was 3D auto-refined with C1 symmetry to 4.3Å after CTF refinement and particle polishing. One round of signal-subtraction and 3D classification using the top mask was performed, giving two major classes, once again, one in the closed conformation and the other in the one-RBD-up conformation. This one-RBD-up class was combined with the RBD-up class from the previous third round of 3D classification, and refined to give a map at 4.6Å resolution from 67,997 particles. The class in the closed state with 63,146 particles was refined to produce a map at 3.8Å resolution after CTF refinement and particle polishing. For data set 2, 3D classification identified two classes in the closed conformation and combination of the two gave a map at 4.9Å resolution from 171,733 particles after 3D auto-refinement. One round of signal-subtraction and 3D classification using the top mask was performed, leading to two major classes, one in the closed conformation and the other in the one-RBD-up conformation. Another round of 3D auto-refinement was carried out, giving two maps with 71,425 and 42,005 particles, respectively.

The two classes in the closed state from the two data sets were then combined and subjected to 3D auto-refinement, CTF refinement, particle polishing and a final round of 3D auto-refinement, yielding a map at 4.1Å resolution from 255,909 particles. One round of signal-subtraction 3D and classification with the top mask was performed, leading to a major class in the closed conformation. After 3D auto-refinement with C1/C3 symmetry using a whole mask, CTF refinement, particle polishing and a final round of auto-refinement, this class yielded a final reconstruction at 3.1Å resolution from 123,193 particles. Similarly, the one-RBD-up classes from the two date sets were combined and subjected to 3D auto-refinement, CTF refinement, particle polishing and 3D auto-refinement, producing a map at 4.0Å resolution from 110,002 particles. One round of signal-subtraction and 3D classification using the top mask was carried out, giving two major classes in distinct one-RBD-up conformations with the RBD projecting at slightly different angles. After 3D auto-refinement with C1 symmetry using a whole mask, CTF refinement, particle polishing and a final round of auto-refinement, the two classes yielded two reconstructions at 3.7Å and 4.3Å resolution from 81,717 and 21,830 particles, respectively. Additional rounds of 3D auto-refinement were performed for each class using different sizes of masks at the top region to improve local resolution. The best density maps were used for model building.

For the Gamma sample, which was not very stable and quite heterogeneous, three independent data sets were collected. 1,652,420, 2,757,190 and 1,893,863 particles were extracted by crYOLO with a trained model from 25,424, 32,569 and 29,163 images, respectively, from the three sets. The selected particles were subjected to 2D classification, giving 1,564,938, 1,904,078 and 1,788,000 particles, respectively. A low-resolution negative-stain reconstruction of the Wuhan-Hu-1 (D614) sample was low-pass-filtered to 40Å as an initial model for 3D classification with C1 symmetry. Two or three rounds of 3D classification were performed separately for the three data sets and each of them produced a major class resembling an S trimer. The three classes were combined and subjected to additional round of 3D classification in C1 symmetry, leading to three good classes. The new classes were auto-refined, giving a map at 4.9Å resolution from 143,872 particles. A round of signal-subtraction and 3D classification using the top mask were performed to produce two major classes in two distinct one-RBD-up conformations with the RBD projecting at slightly different angles. The two classes contain 69,302 and 36,346 particles, respectively and they were subjected to 3D auto-refinement with C1 symmetry using a whole mask, CTF refinement/particle polishing and a final round auto-refinement, yielding two reconstructions at 3.8Å and 4.4Å resolution, respectively. Additional rounds of 3D auto-refinement were performed for each class using different sizes of masks at the top region to improve local resolution. The best density maps were used for model building.

All resolutions were reported from the gold-standard Fourier shell correlation (FSC) using the 0.143 criterion. Density maps were corrected from the modulation transfer function of the K3 detector and sharpened by applying a temperature factor that was estimated using post-processing in RELION. Local resolution was determined using ResMap (*56*) with half-reconstructions as input maps.

### Model building

The initial templates for model building used our G614 S trimer structures (PDB ID: 7KRQ and PDB ID: 7KRR; ref(*31*)). Several rounds of manual building were performed in Coot (*57*). The model was then refined in Phenix (*58*) against the 3.1Å (closed), 3.4Å (one-RBD-up 1), 4.3Å (one-RBD-up 2) cryo-EM maps of the Delta variant; the 3.8Å (one-RBD-up 1) and 4.4Å (one-RBD-up 2) cryo-EM maps of the Gamma variant; and the 3.1Å (closed), 3.7Å (one-RBD-up 1) and 4.3Å (one-RBD-up 2) cryo-EM maps of the Kappa variant. Iteratively, refinement was performed in both Refmac (real space refinement) and ISOLDE (*59*), and the Phenix refinement strategy included minimization_global, local_grid_search, and adp, with rotamer, Ramachandran, and reference-model restraints, using 7KRQ and 7KRR as the reference model. The refinement statistics are summarized in Table S3. Structural biology applications used in this project were compiled and configured by SBGrid (*60*).

## Acknowledgments

We thank the SBGrid team for technical assistance, K. Arnett for support and advice on the BLI experiments, and S. Harrison and A. Carfi for critical reading of the manuscript. EM data were collected at the Harvard Cryo-EM Center for Structural Biology of Harvard Medical School. We acknowledge support for COVID-19 related structural biology research at Harvard from the Nancy Lurie Marks Family Foundation and the Massachusetts Consortium on Pathogen Readiness (MassCPR). This work was supported by Fast grants by Emergent Ventures (to B.C. and D.R.W.), COVID-19 Award by Massachusetts Consortium on Pathogen Readiness (MassCPR; to B.C. and D.R.W.), and NIH grants AI147884 (to B.C.), AI141002 (to B.C.), AI127193 (to B.C. and James Chou) and AI39538 (to D.R.W).

## Author Contribution

B.C., J.Z., T.X. and Y.C. conceived the project. Y.C. expressed and purified the full-length S proteins with help from H.P., and carried out negative stain EM. T.X. performed BLI and cell-cell fusion experiments. J.Z. prepared cryo grids and performed EM data collection with contributions from M.L.M. and R.M.W., and processed the cryo-EM data, built and refined the atomic models. J.L. created all the expression constructs and performed the neutralization assays using the MLV-based pseudoviruses. C.L.L. and M.S.S performed the neutralization assays using the HIV-based pseudoviruses. H.Z., K.A. and W.Y. performed the flow cytometry experiments. P.T., A.G. and, D.R.W. produced anti-S monoclonal antibodies. S.R.V. contributed to cell culture and protein production. All authors analyzed the data. B.C., J.Z., T.X. and Y.C. wrote the manuscript with input from all other authors.

