## Supplemental Figures and Tables for "Membrane fusion and immune evasion by the spike protein of SARS-CoV-2 Delta variant"

### Supplementary materials

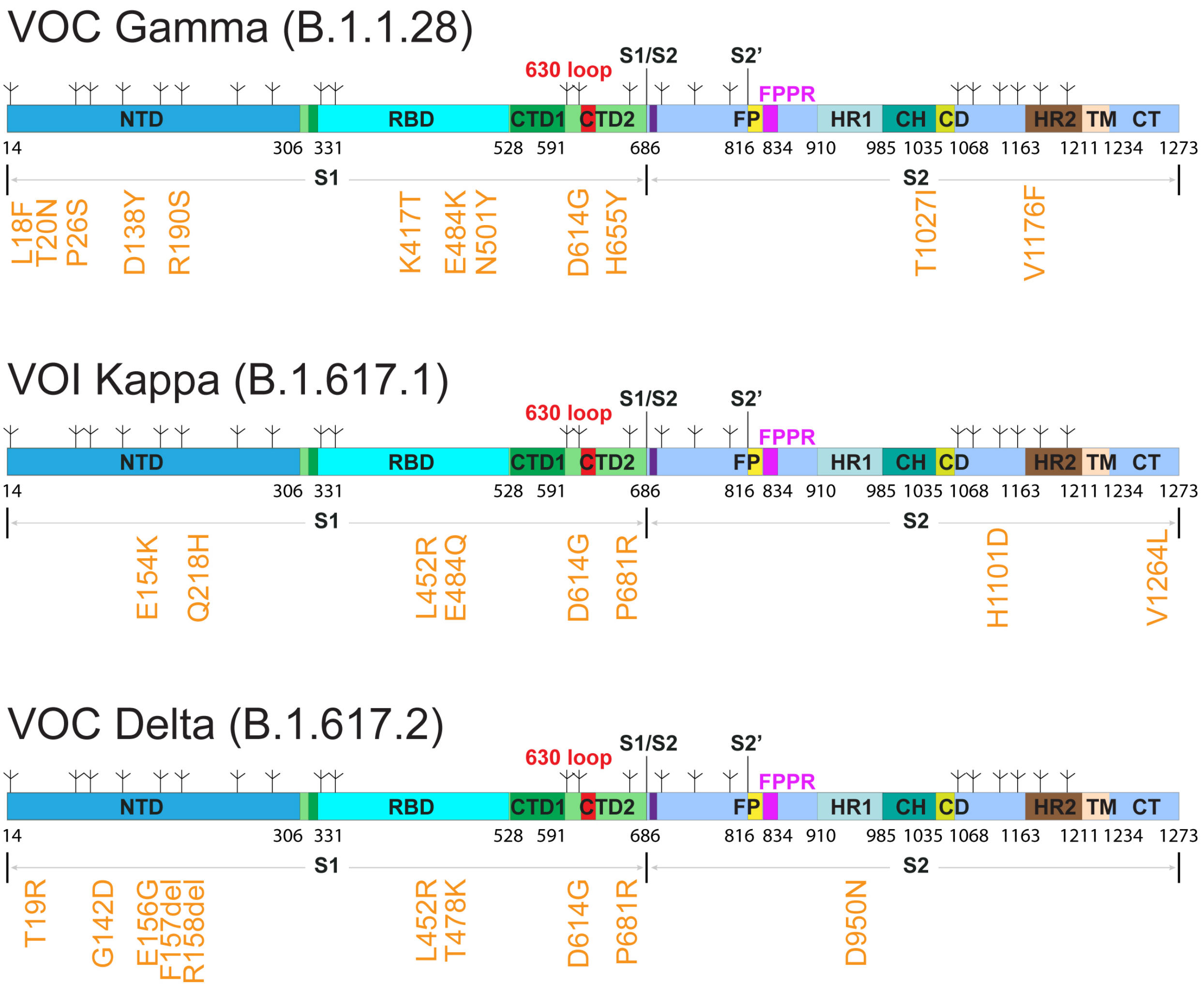


**Figure S1. Schematic representation of full-length SARS-CoV-2 spike (S) proteins from the VOC Gamma (B.1.1.28), VOI Kappa (B.1.617.1) and VOC Delta (B.1.617.2).** The sequences are derived from the Gamma (hCoV-19/Brazil/AM-992/2020), Kappa (hCoV-19/India/MH-NEERI-NGP-40449/2021) and Delta (hCoV-19/India/GJ-GBRC619/2021) variants. Segments of S1 and S2 include: NTD, N-terminal domain; RBD, receptor-binding domain; CTD1, C-terminal domain 1; CTD2, C-terminal domain 2; 630 loop; S1/S2, S1/S2 cleavage site; S2', S2' cleavage site; FP, fusion peptide; FPPR, fusion peptide proximal region; HR1, heptad repeat 1; CH, central helix region; CD, connector domain; HR2, heptad repeat 2; TM, transmembrane anchor; CT, cytoplasmic tail; and tree-like symbols for glycans. Positions of all mutations (from the amino-acid sequence of Wuhan-Hu-1) are shown in orange text.

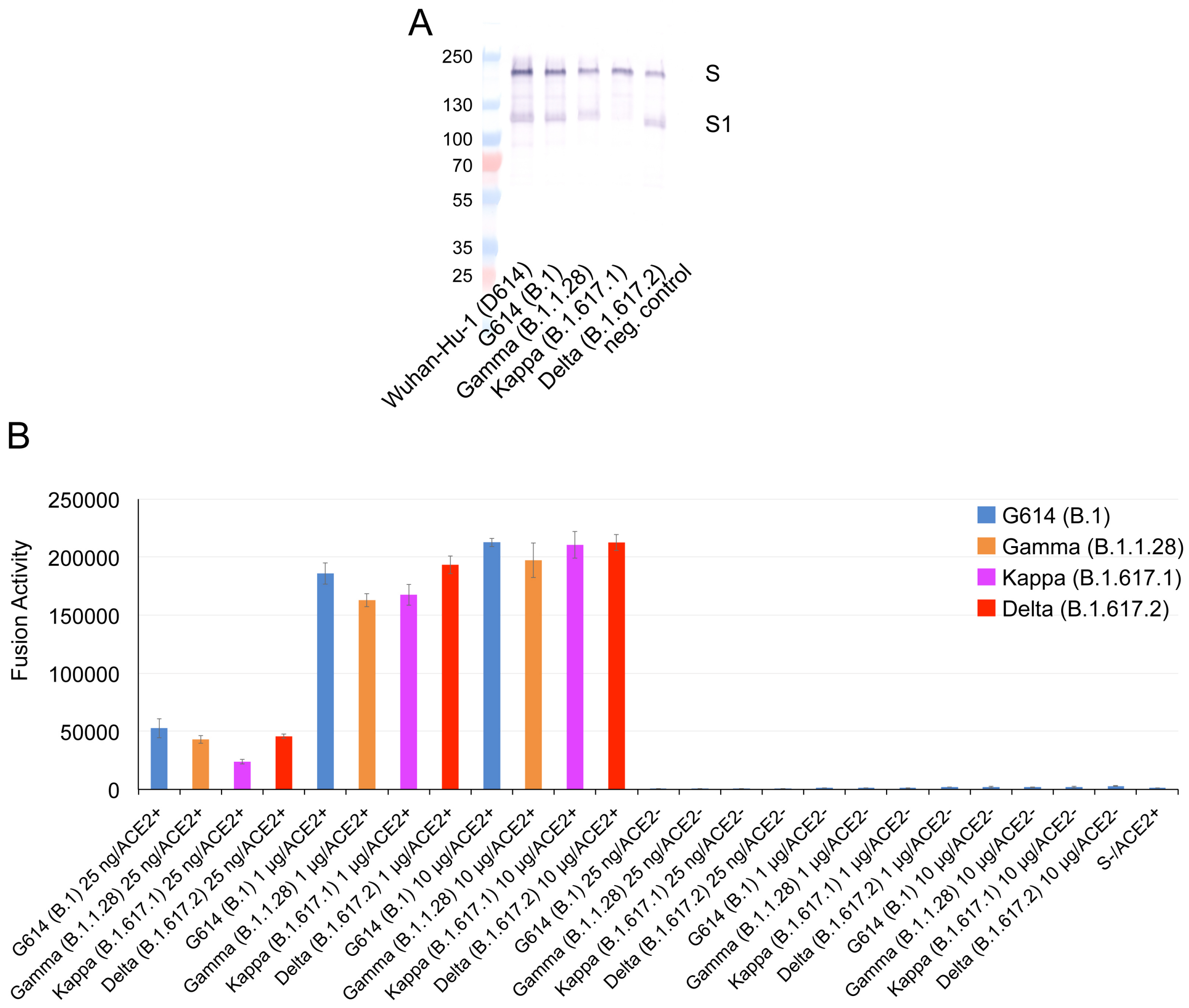


**Figure S2. Expression and cell-cell fusion of SARS-CoV-2 variant S proteins. (A)** Expression and processing of the full-length S constructs in HEK293 cells. S samples prepared from HEK293 cells transiently transfected with 10 μg of the full-length S expression plasmids were detected by anti-RBD polyclonal antibodies. Bands for the uncleaved S and S1 fragment are indicated. (**B**) HEK293T cells transfected with the untagged, full-length S protein expression plasmids were fused with ACE2-expressing cells. Cell-cell fusion led to reconstitution of α and ω fragments of β-galactosidase yielding an active enzyme, and the fusion activity was then quantified by a chemiluminescent assay. No ACE2 and no S were negative controls.

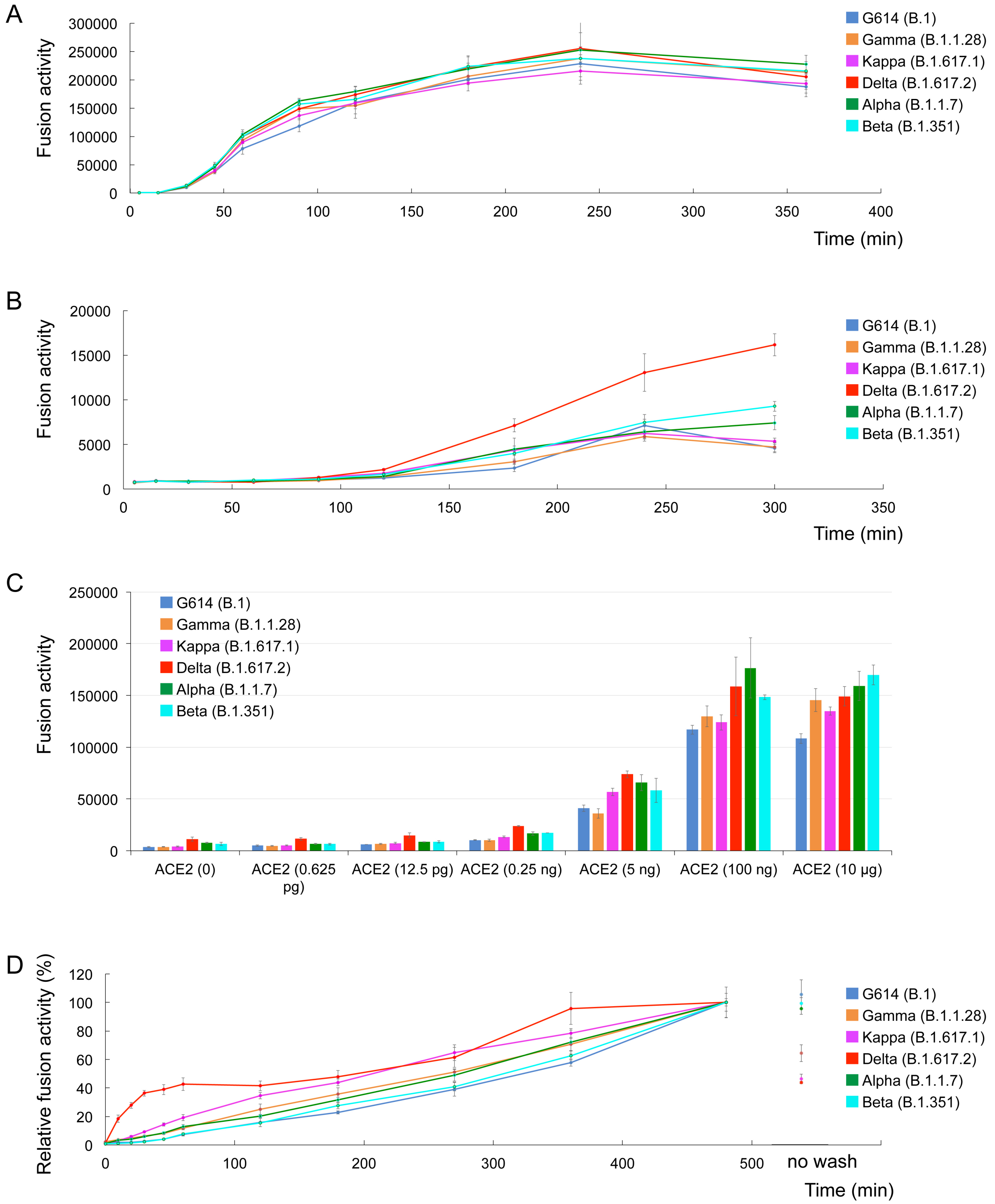


**Figure S3. Comparison of membrane fusion mediated by the Delta variant with that by other variants.** (**A**) Time-course of cell-cell fusion mediated by various full-length S proteins expressed at the 10 μg transfection level, as indicated, with HEK293 cells transfected with 10 μg of ACE2. (**B**) Time-course of cell-cell fusion mediated by various full-length S proteins (10 μg transfection), as indicated, with HEK293 cells with no exogenous ACE2. (**C**) Cell-cell fusion mediated by various full-length S proteins with HEK293 cells transfected with various levels (0-10 μg) of ACE2 expression constructs. (**D**) Time course of infection HEK293-ACE2 cells by MLV-based pseudotyped viruses using various SARS-CoV-2 variant S constructs containing a CT deletion in a single cycle. Infection was initiated by mixing viruses and target cells, and viruses were washed out at each time point as indicated. Selected data points are highlighted in Fig. 1. The experiments were repeated at least three times with independent samples giving similar results.

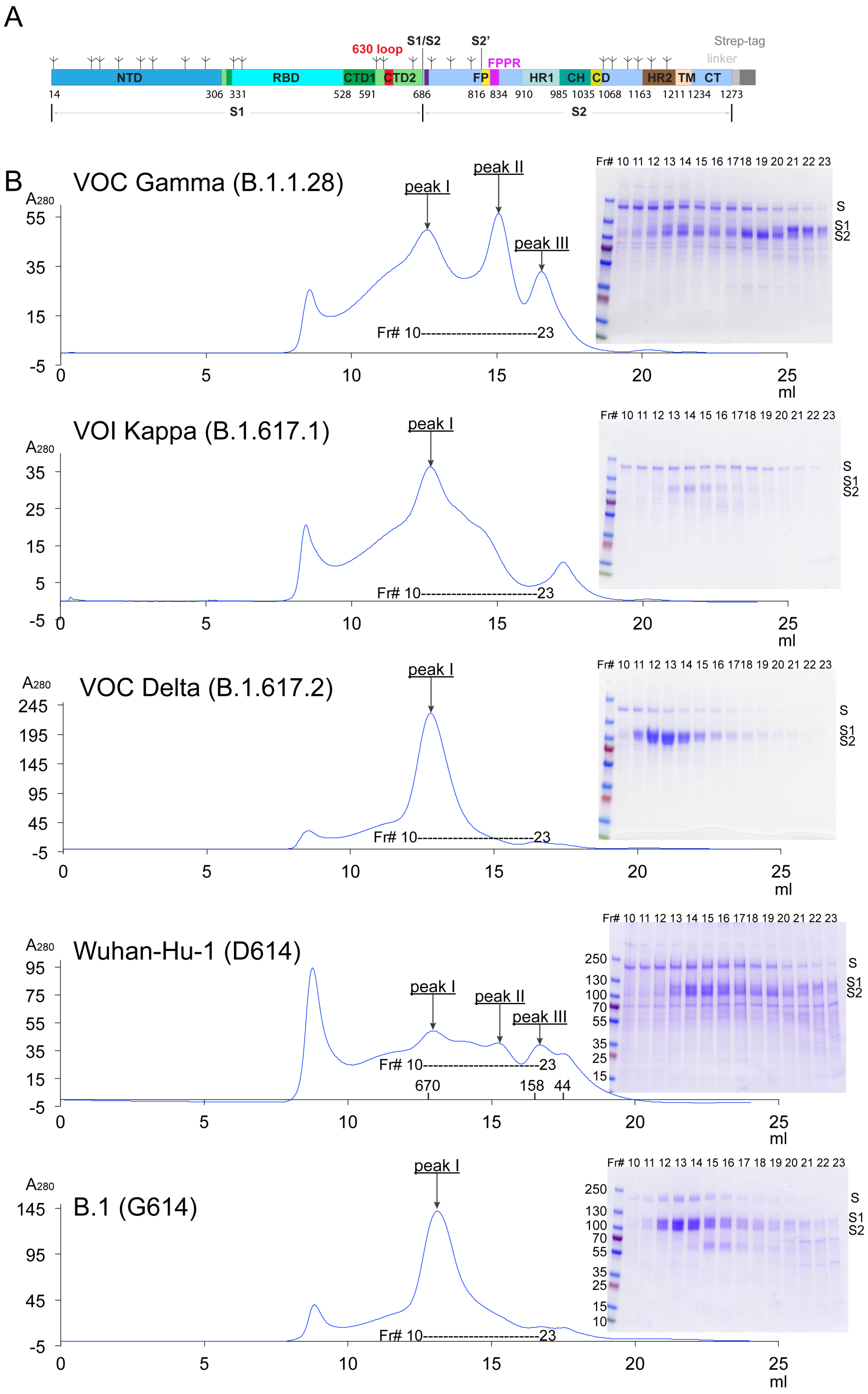

A
630 loop
S1/S2
S2'
Strep-tag
linker
FPPR
NTD
RBD
CTD1
CTD2
FP
HR1
CH
CD
HR2
TM
CT
14 306 331 528 591 686 816 834 910 985 1035 1068 1163 1211 1234 1273
S1
S2
B
VOC Gamma (B.1.1.28)
peak I
peak II
peak III
Fr# 10-----------------------23
VOI Kappa (B.1.617.1)
VOC Delta (B.1.617.2)
Wuhan-Hu-1 (D614)
670 158 44
B.1 (G614)
A280
ml
Fr# 10 11 12 13 14 15 16 1718 19 20 21 22 23
S
S1
S2
250 130 100 70 55 35 25 15 10

**Figure S4. Production of full-length S protein from the Gamma, Kappa and Delta variants.** (**A**) A strep-tag was fused to the C-terminus of the full-length S protein by a flexible linker. (**B**) The full-length S proteins were extracted and purified in detergent DDM, and further resolved by gel-filtration chromatography on a Superose 6 column. Peak I, the prefusion S trimer; peak II, the postfusion S2 trimer; and peak III, the dissociated monomeric S1. Inset, peak fractions were analyzed by Coomassie stained SDS-PAGE. Labeled bands are S, S1 and S2. Fr#, fraction number. Each experiment was repeated at least three times independently with similar results. The data for the preparations from the Wuhan-Hu-1 (D614) and B.1 (G614), published previously (*28, 31*), are included for convenient comparison.

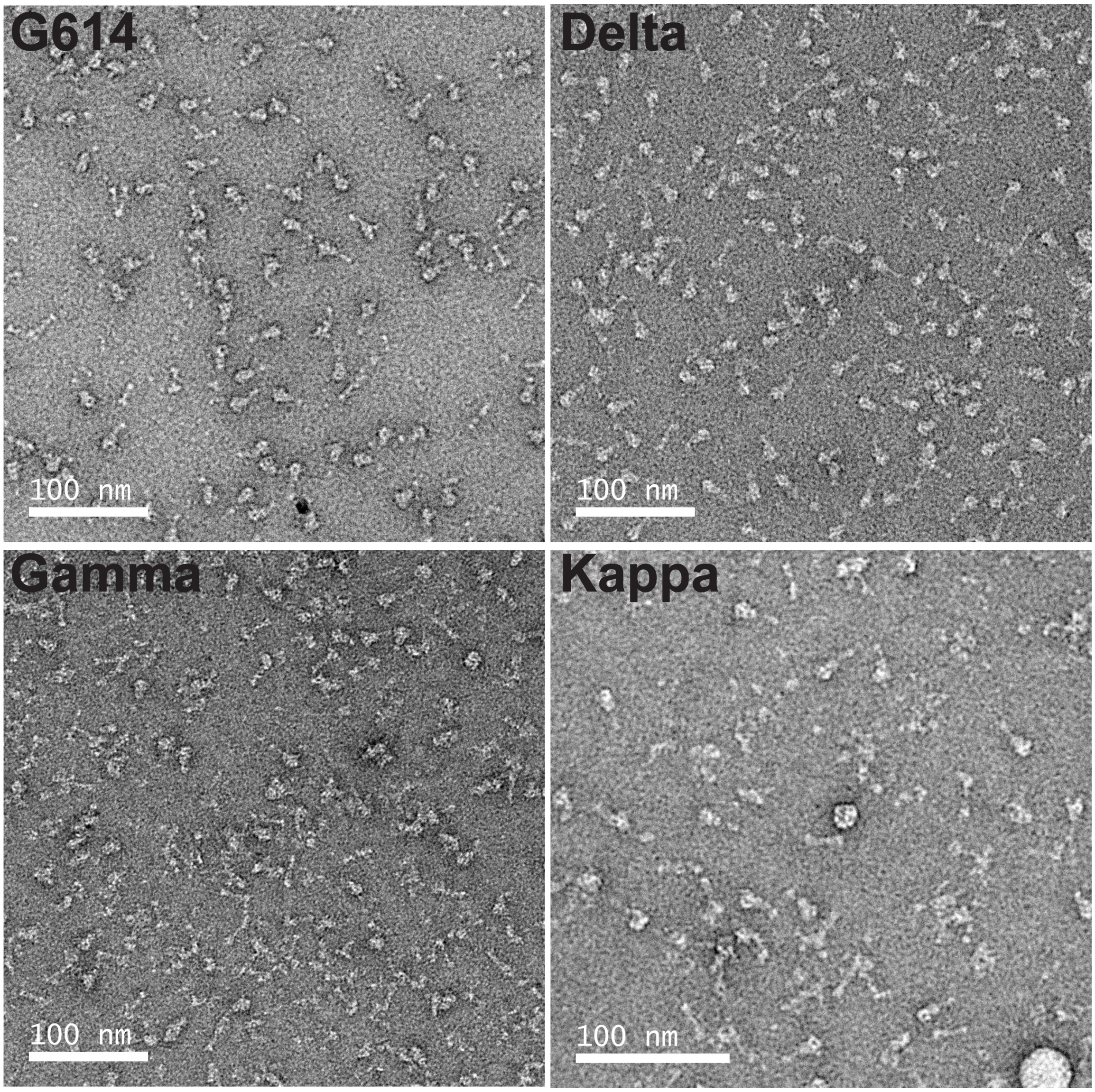


**Figure S5. Representative images of various full-length S trimers by negative stain EM.**

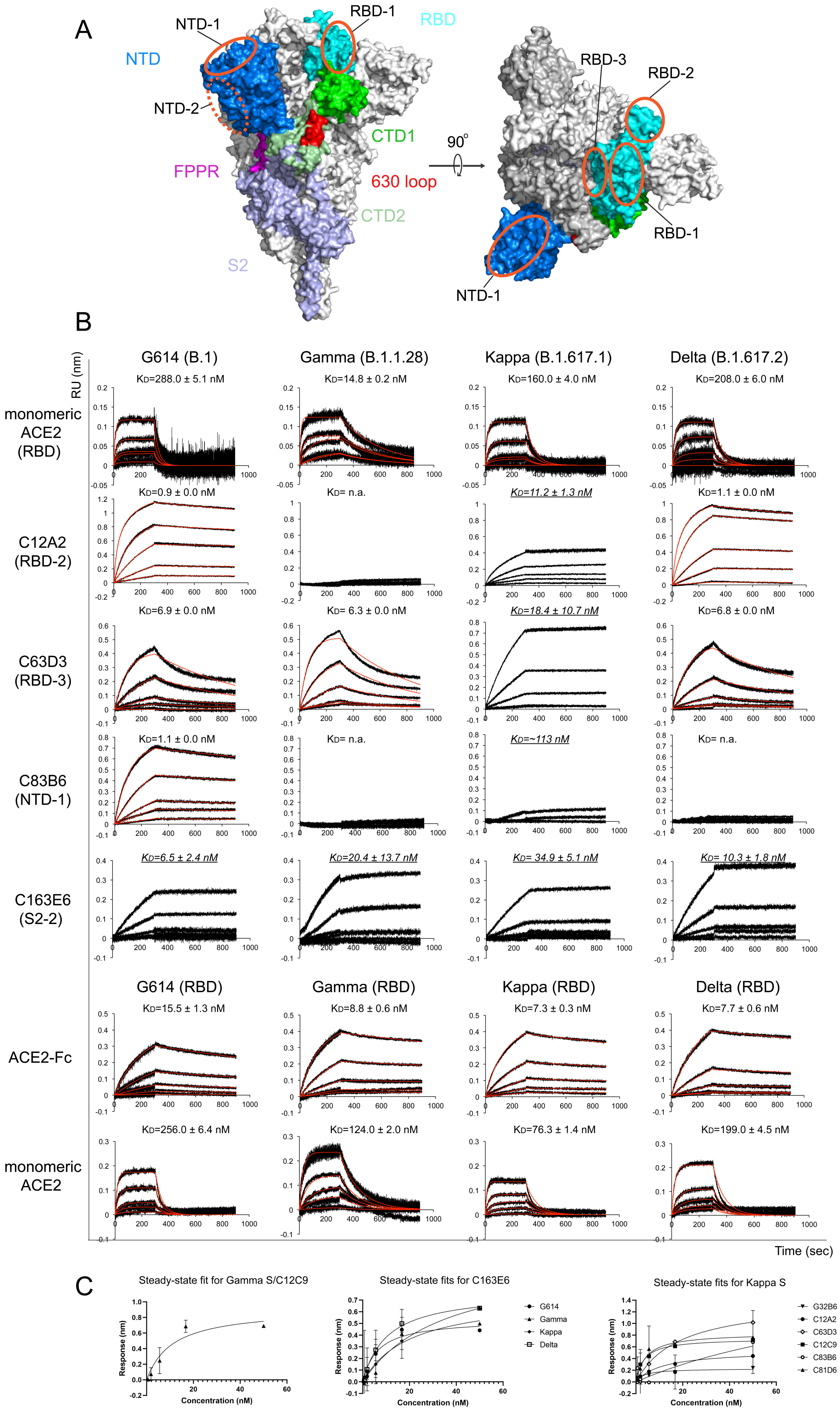

A
NTD-1
RBD-1
RBD
NTD
NTD-2
CTD1
FPPR
630 loop
CTD2
S2
90°
RBD-3
RBD-2
RBD-1
NTD-1
B
G614 (B.1)
Gamma (B.1.1.28)
Kappa (B.1.617.1)
Delta (B.1.617.2)
RU (nm)
monomeric ACE2 (RBD)
KD=288.0 ± 5.1 nM
KD=14.8 ± 0.2 nM
KD=160.0 ± 4.0 nM
KD=208.0 ± 6.0 nM
C12A2 (RBD-2)
KD=0.9 ± 0.0 nM
KD= n.a.
KD=11.2 ± 1.3 nM
KD=1.1 ± 0.0 nM
C63D3 (RBD-3)
KD=6.9 ± 0.0 nM
KD= 6.3 ± 0.0 nM
KD=18.4 ± 10.7 nM
KD=6.8 ± 0.0 nM
C83B6 (NTD-1)
KD=1.1 ± 0.0 nM
KD= n.a.
KD=~113 nM
KD= n.a.
C163E6 (S2-2)
KD=6.5 ± 2.4 nM
KD=20.4 ± 13.7 nM
KD= 34.9 ± 5.1 nM
KD= 10.3 ± 1.8 nM
G614 (RBD)
Gamma (RBD)
Kappa (RBD)
Delta (RBD)
ACE2-Fc
KD=15.5 ± 1.3 nM
KD=8.8 ± 0.6 nM
KD=7.3 ± 0.3 nM
KD=7.7 ± 0.6 nM
monomeric ACE2
KD=256.0 ± 6.4 nM
KD=124.0 ± 2.0 nM
KD=76.3 ± 1.4 nM
KD=199.0 ± 4.5 nM
Time (sec)
C
Steady-state fit for Gamma S/C12C9
Steady-state fits for C163E6
Steady-state fits for Kappa S
Response (nm)
Concentration (nM)
G614
Gamma
Kappa
Delta
G32B6
C12A2
C63D3
C12C9
C83B6
C81D6

**Figure S6. Additional antigenic properties of the purified full-length SARS-CoV-2 S proteins**. (**A**) Antibody competition clusters as described in ref(*32*). Surface regions of the S trimer targeted by antibodies on S1 are highlighted by orange ellipses, including RBD-1, RBD-2, RBD-3, NTD-1 and NTD-2. The exact location of NTD-2 is uncertain and therefore marked with a dashed line. (**B**) Binding analysis of the prefusion S trimers from G614, Gamma, Kappa and Delta variants with soluble ACE2 constructs was performed by bio-layer interferometry (BLI). For ACE2 binding, the purified S proteins were immobilized on AR2G biosensors and dipped into the wells containing ACE2 at various concentrations. For antibody binding, various antibodies were immobilized to AHC biosensors and dipped into the wells containing each purified S protein at different concentrations. Binding kinetics were evaluated using a 1:1 Langmuir model except for antibody C12A2 targeting the RBD-2, which was analyzed by a bivalent binding model. The sensorgrams are in black and the fits in red. Binding constants highlighted by underlines were estimated by steady-state analysis as described in the Methods. RU, response unit. Binding constants are also summarized here and in Table S1. All experiments were repeated at least twice with essentially identical results. (**C**) Steady-state analysis by plotting steady-state responses against concentrations. Kd values were derived from the fits.

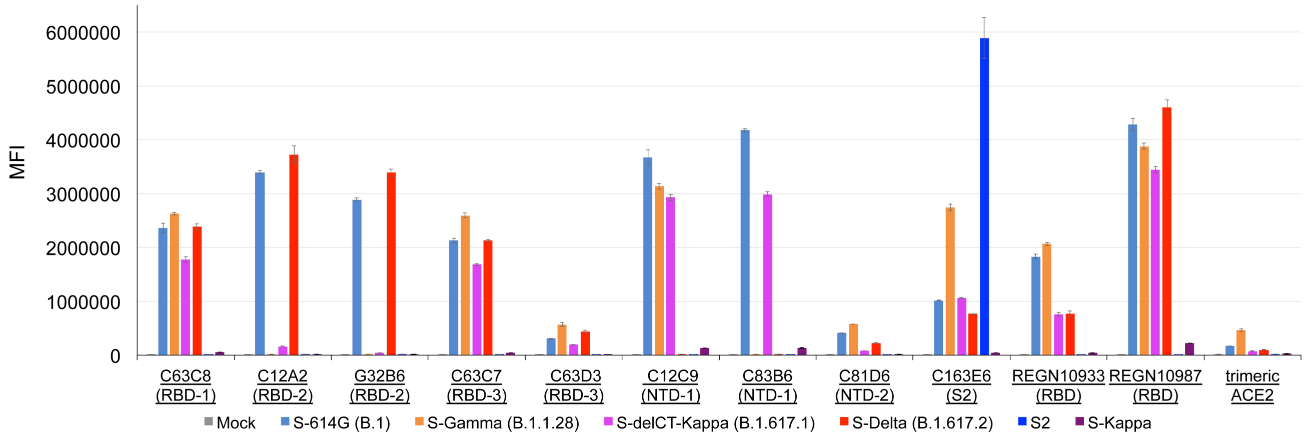


**Figure S7. Antigenic properties of the cell-surface S proteins assessed by flow cytometry.** Antibody and ACE2 binding to the full-length S proteins of the G614, Gamma, Kappa and Delta variants, as well as an S2 construct expressed on the cell surfaces analyzed by flow cytometry. An S-delCT-Kappa with the C-terminal 19 residues deleted was included because the expression level of the full-length S construct (S-Kappa) was low. The antibodies and their targets are indicated. Two therapeutic anti-RBD antibodies by Regeneron, REGN10933 (casirivimab) and REGN10987 (imdevimab), were also included in this assay. A designed ACE2-based inhibitor $ACE2_{615}$-foldon-T27W was used for detecting receptor binding (*33*). MFI, mean fluorescent intensity. The error bars represent standard errors of mean from measurements using three independently transfected cell samples. The flow cytometry assays were repeated three times with essentially identical results.

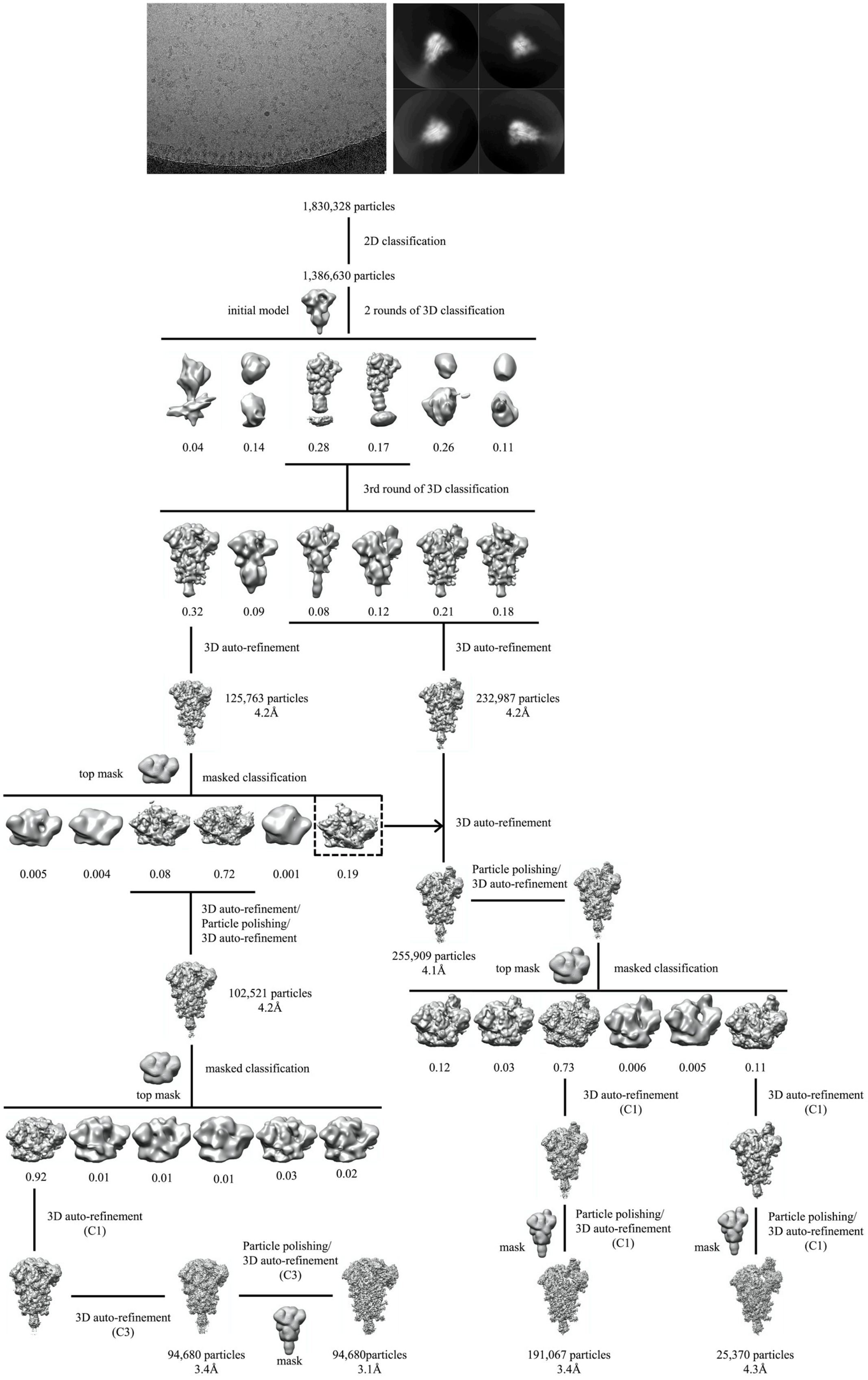

1,830,328 particles
2D classification
1,386,630 particles
initial model
2 rounds of 3D classification
0.04
0.14
0.28
0.17
0.26
0.11
3rd round of 3D classification
0.32
0.09
0.08
0.12
0.21
0.18
3D auto-refinement
3D auto-refinement
125,763 particles
4.2Å
232,987 particles
4.2Å
top mask
masked classification
3D auto-refinement
0.005
0.004
0.08
0.72
0.001
0.19
Particle polishing/
3D auto-refinement
3D auto-refinement/
Particle polishing/
3D auto-refinement
255,909 particles
4.1Å
top mask
masked classification
102,521 particles
4.2Å
0.12
0.03
0.73
0.006
0.005
0.11
masked classification
top mask
3D auto-refinement
(C1)
3D auto-refinement
(C1)
0.92
0.01
0.01
0.01
0.03
0.02
3D auto-refinement
(C1)
mask
Particle polishing/
3D auto-refinement
(C1)
mask
Particle polishing/
3D auto-refinement
(C1)
Particle polishing/
3D auto-refinement
(C3)
3D auto-refinement
(C3)
94,680 particles
3.4Å
mask
94,680particles
3.1Å
191,067 particles
3.4Å
25,370 particles
4.3Å

**Figure S8. Cryo-EM analysis of the Delta S trimer.** Top, representative micrograph**,** and 2D averages (box dimension: 396Å) of the cryo-EM particle images of the Delta S trimer. Bottom, data processing workflow for structure determination.

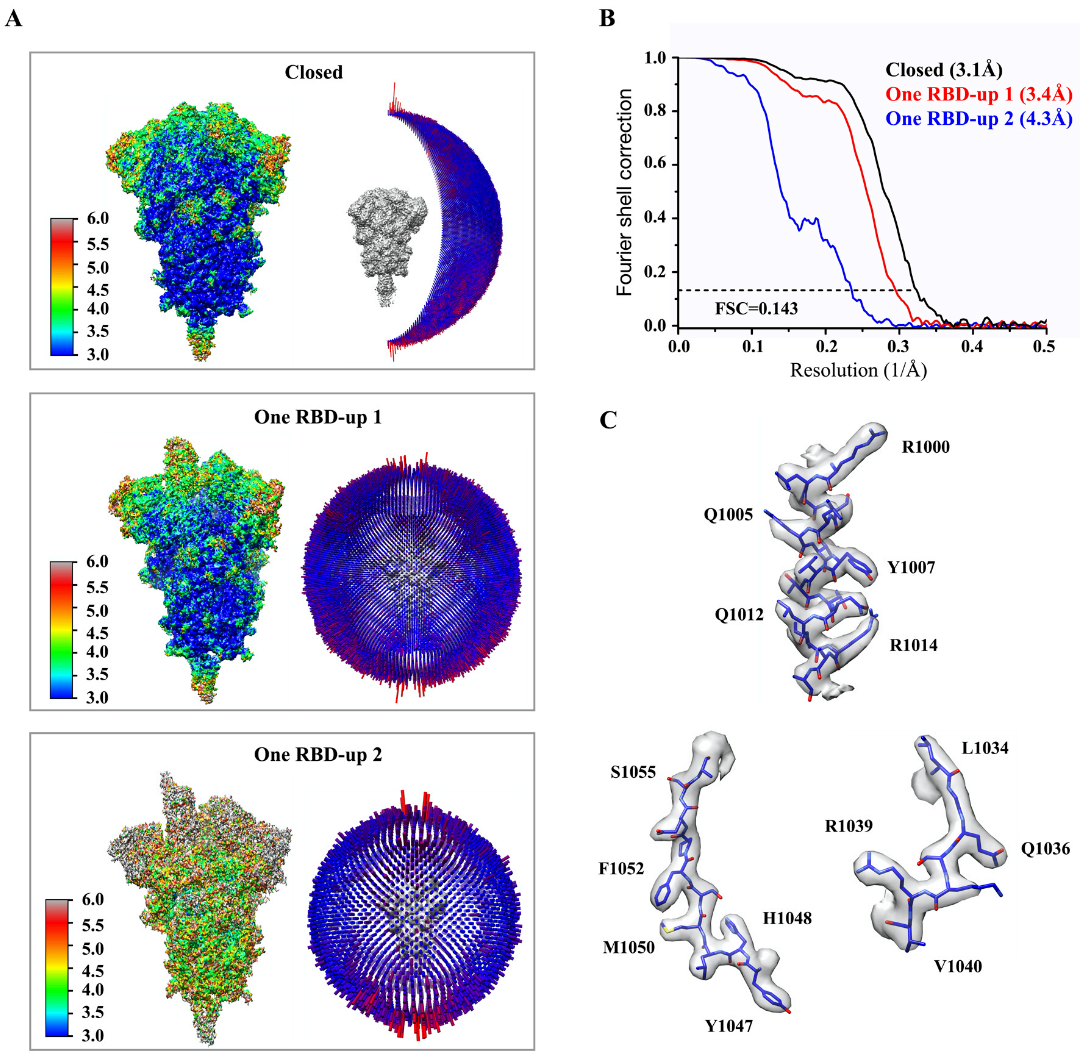


**Figure S9. Analysis of the Delta S trimer structure.** (**A**) 3D reconstructions of the Delta trimer preparation in the closed, two one-RBD-up conformations, respectively, are colored according to local resolution estimated by ResMap. Angular distribution of the cryo-EM particles used in each reconstruction is shown in the side view of the EM map. (**B**) Gold standard FSC curves of the three refined 3D reconstructions of the Delta S trimer. (**C**) Representative density in gray surface from EM maps with a resolution better than 3.5Å.

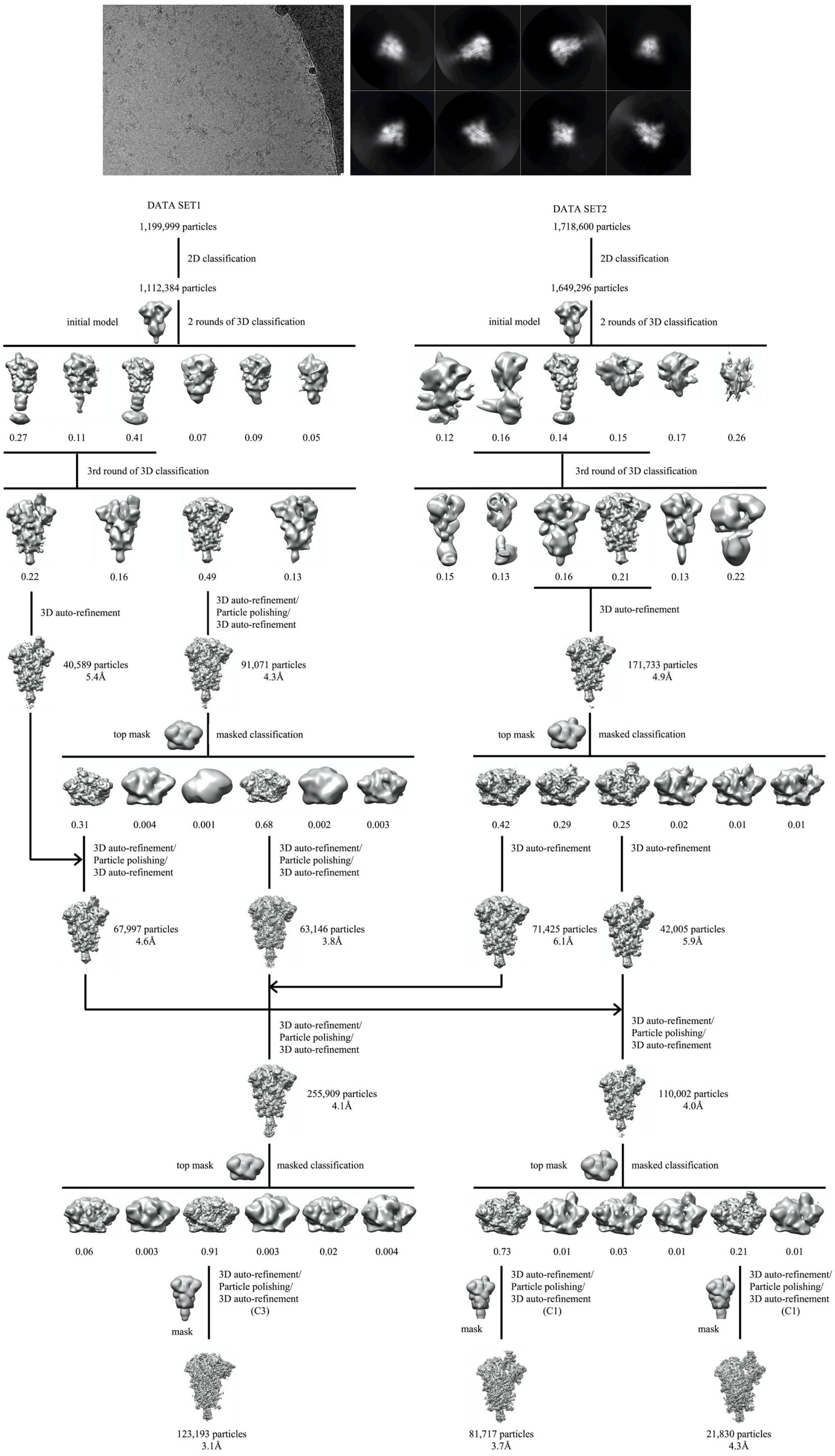

DATA SET1
1,199,999 particles
2D classification
1,112,384 particles
initial model
2 rounds of 3D classification
0.27
0.11
0.41
0.07
0.09
0.05
3rd round of 3D classification
0.22
0.16
0.49
0.13
3D auto-refinement
3D auto-refinement/
Particle polishing/
3D auto-refinement
40,589 particles
5.4Å
91,071 particles
4.3Å
top mask
masked classification
0.31
0.004
0.001
0.68
0.002
0.003
3D auto-refinement/
Particle polishing/
3D auto-refinement
3D auto-refinement/
Particle polishing/
3D auto-refinement
67,997 particles
4.6Å
63,146 particles
3.8Å
DATA SET2
1,718,600 particles
2D classification
1,649,296 particles
initial model
2 rounds of 3D classification
0.12
0.16
0.14
0.15
0.17
0.26
3rd round of 3D classification
0.15
0.13
0.16
0.21
0.13
0.22
3D auto-refinement
171,733 particles
4.9Å
top mask
masked classification
0.42
0.29
0.25
0.02
0.01
0.01
3D auto-refinement
3D auto-refinement
71,425 particles
6.1Å
42,005 particles
5.9Å
3D auto-refinement/
Particle polishing/
3D auto-refinement
3D auto-refinement/
Particle polishing/
3D auto-refinement
255,909 particles
4.1Å
110,002 particles
4.0Å
top mask
masked classification
0.06
0.003
0.91
0.003
0.02
0.004
top mask
masked classification
0.73
0.01
0.03
0.01
0.21
0.01
mask
3D auto-refinement/
Particle polishing/
3D auto-refinement
(C3)
mask
3D auto-refinement/
Particle polishing/
3D auto-refinement
(C1)
mask
3D auto-refinement/
Particle polishing/
3D auto-refinement
(C1)
123,193 particles
3.1Å
81,717 particles
3.7Å
21,830 particles
4.3Å

**Figure S10. Cryo-EM analysis of the Kappa S trimer.** Top, representative micrograph**,** and 2D averages (box dimension: 396Å) of the cryo-EM particle images of the Kappa S trimer. Bottom, data processing workflow for structure determination.

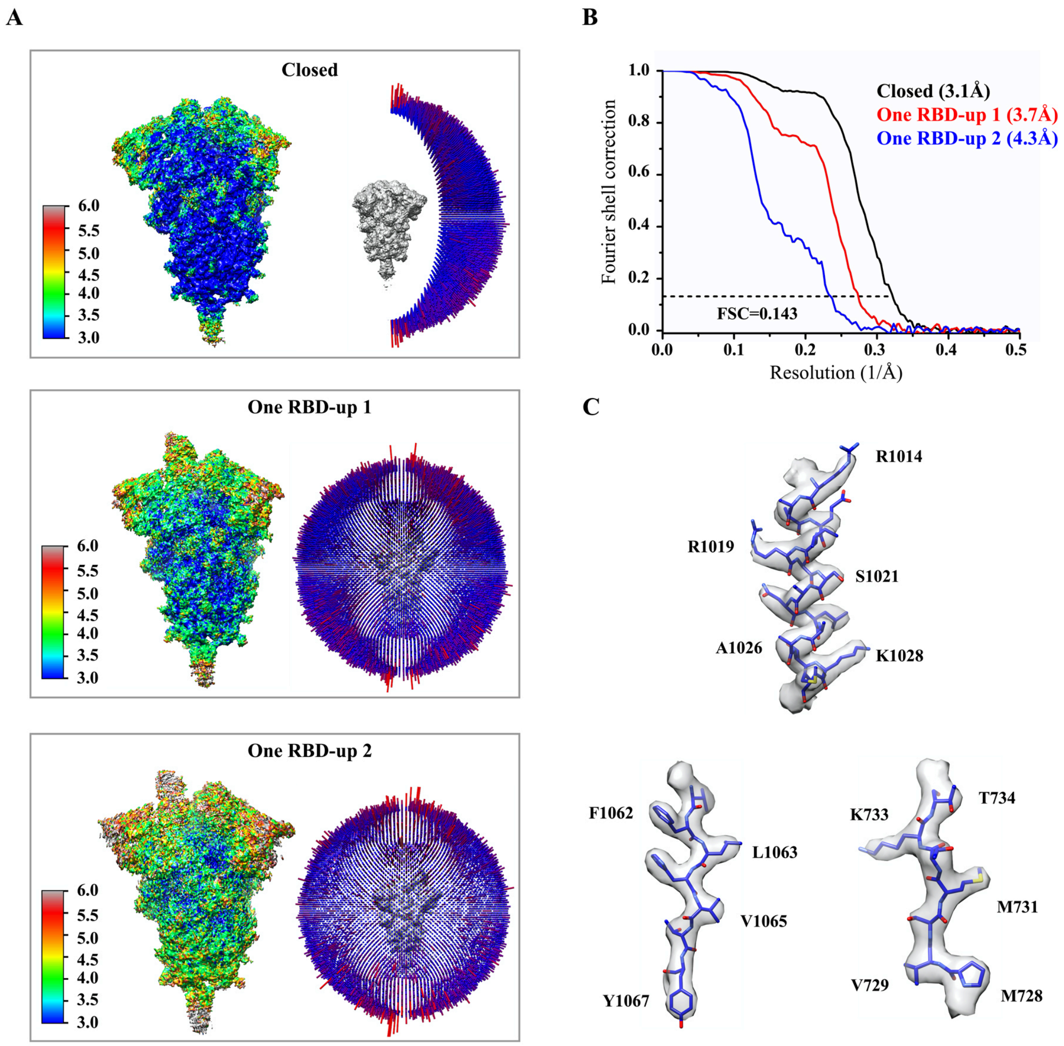


**Figure S11. Analysis of the Kappa S trimer structure.** (**A**) 3D reconstructions of the Kappa S trimer preparation in the closed and two one-RBD-up conformations, respectively, are colored according to local resolution estimated by ResMap. Angular distribution of the cryo-EM particles used in reconstruction for the closed conformation is shown in the side view of the EM map. (**B**) Gold standard FSC curves of the refined 3D reconstructions of the Kappa S trimer. (**C**) Representative density in gray surface from EM maps with a resolution better than 3.5Å.

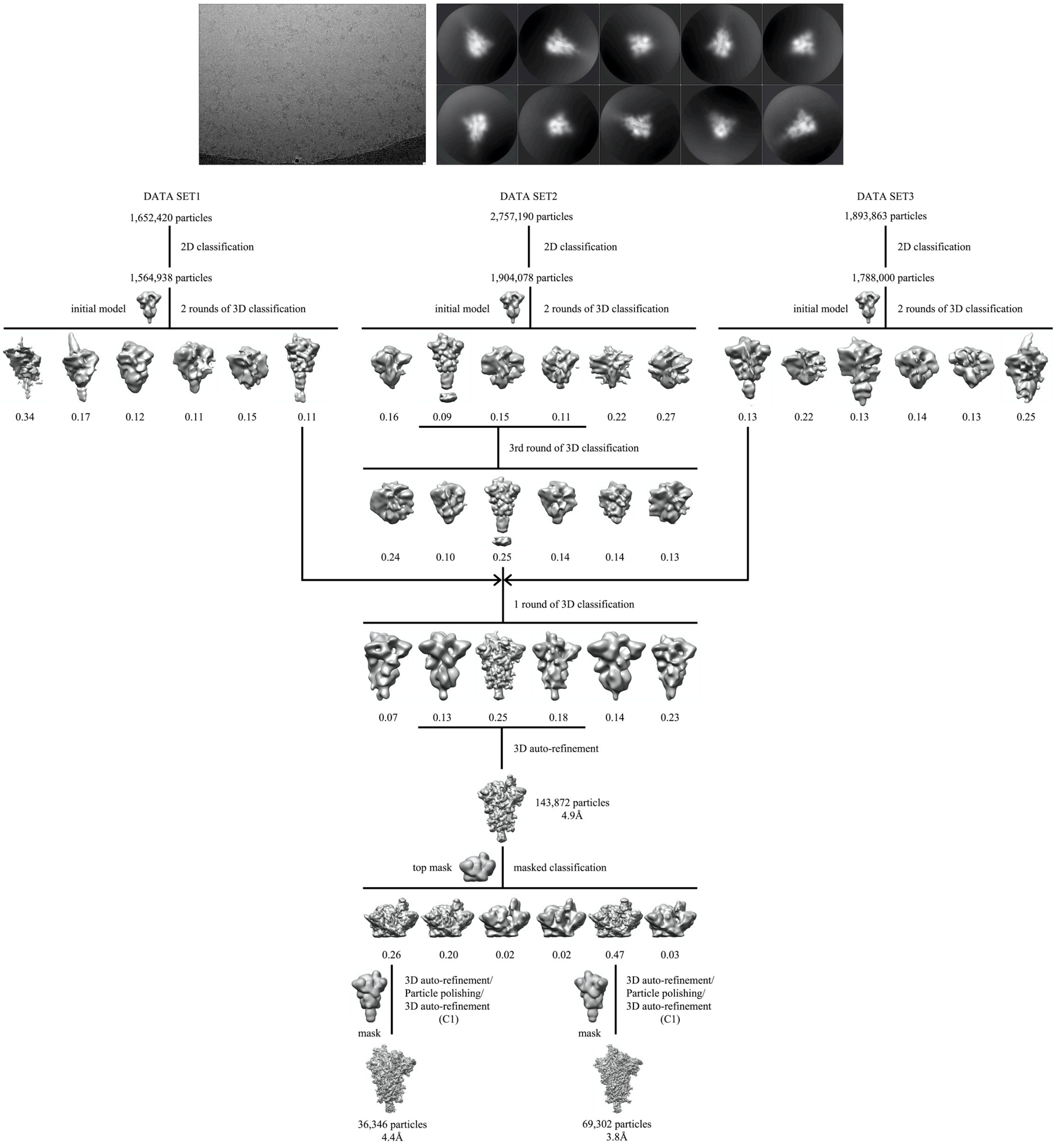


**Figure S12. Cryo-EM analysis of the Gamma S trimer.** Top, representative micrograph**,** and 2D averages (box dimension: 396Å) of the cryo-EM particle images of the Gamma S trimer. Bottom, data processing workflow for structure determination.

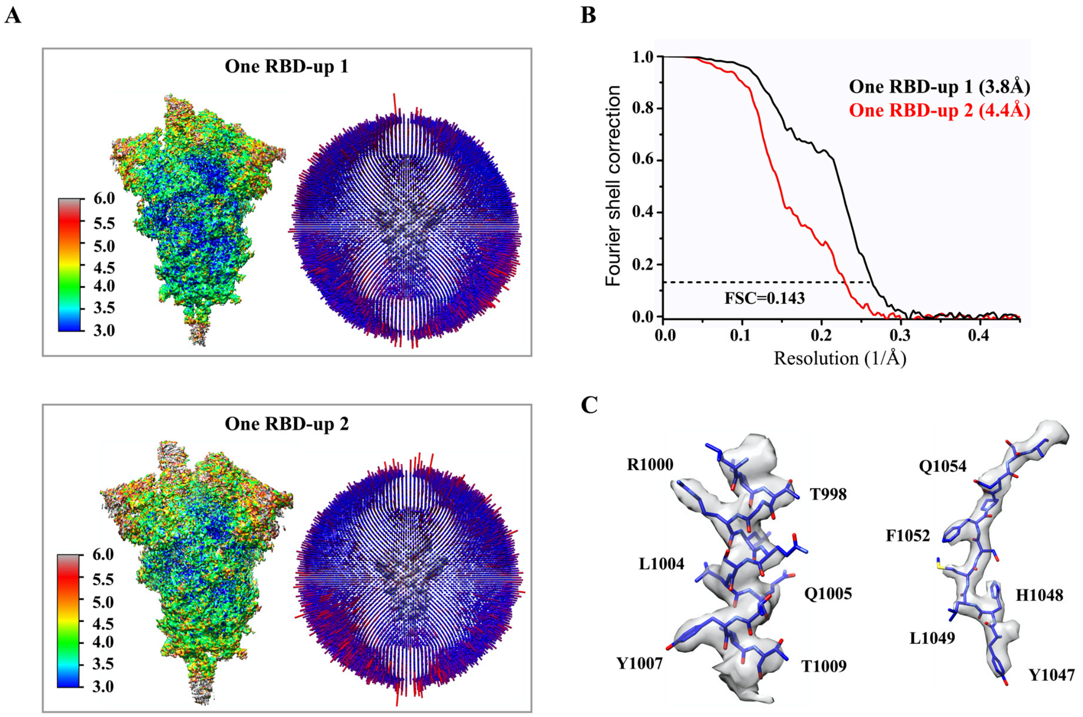


**Figure S13. Analysis of the Gamma S trimer structure.** (**A**) 3D reconstructions of the Gamma S trimer preparation in the two one-RBD-up conformations, respectively, are colored according to local resolution estimated by ResMap. Angular distribution of the cryo-EM particles used in reconstruction for the closed conformation is shown in the side view of the EM map. (**B**) Gold standard FSC curves of the refined 3D reconstructions of the Gamma S trimer. (**C**) Representative density in gray surface from the 3.8Å EM map.

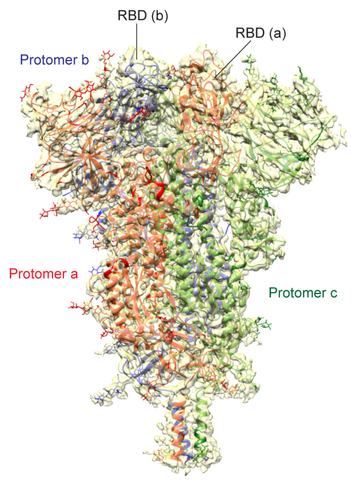

Delta closed conformation
(31%; 3.1Å)

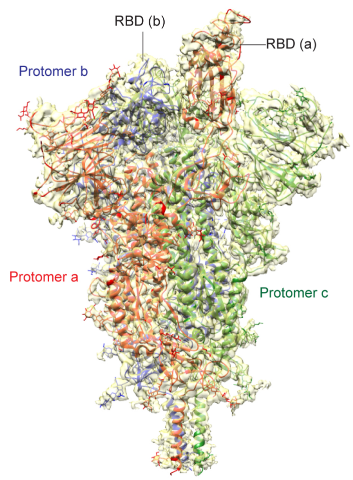

Delta one-RBD-up conformation 1
(61%; 3.4Å)

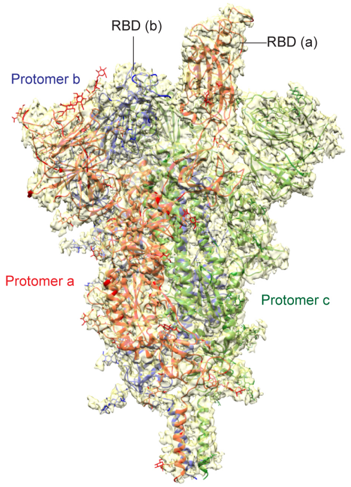

Delta one-RBD-up conformation 2
(8%; 4.3Å)

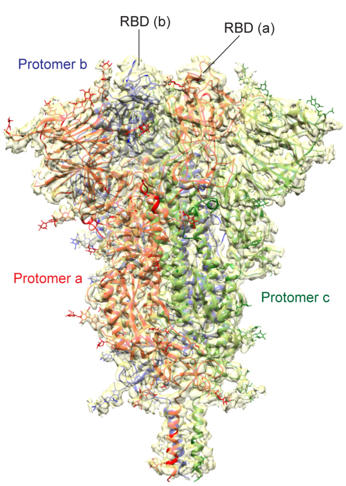

Kappa closed conformation
(54%; 3.1Å)

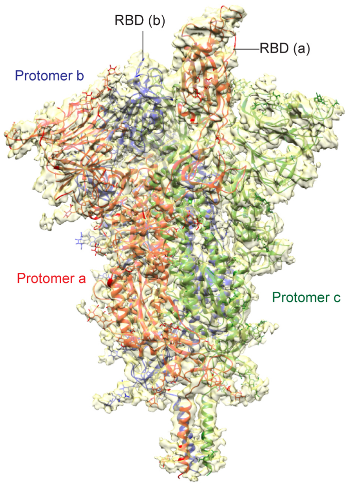

Kappa one-RBD-up conformation 1
(36%; 3.7Å)

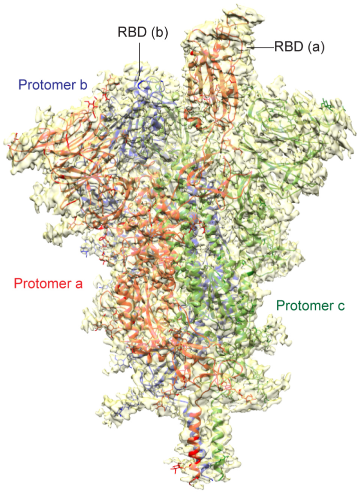

Kappa one-RBD-up conformation 2
(10%; 4.3Å)

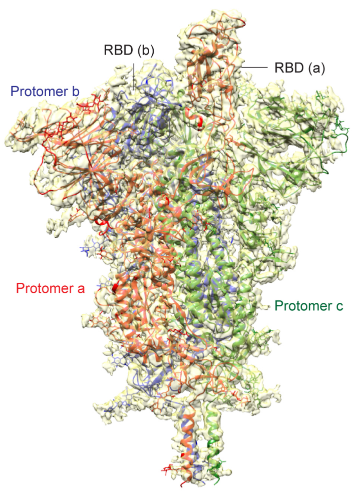

Gamma one-RBD-up conformation 1
(66%; 3.8Å)

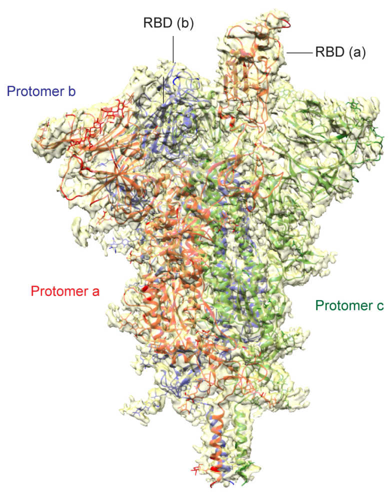

Gamma one-RBD-up conformation 2
(34%; 4.4Å)

**Figure S14. Cryo-EM structures of the full-length S proteins of the Delta, Kappa and Gamma variants**. Three structures of the Delta S trimer, representing a closed prefusion conformation and two distinct one-RBD-up conformations, were modeled based on corresponding cryo-EM density maps at 3.1-4.3Å resolution. Three structures of the Kappa S trimer, representing a closed prefusion conformation and two distinct one-RBD-up conformations, were modeled based on corresponding cryo-EM density maps at 3.1-4.3Å resolution. Two structures of the Gamma S trimer, representing two distinct one-RBD-up conformations, were modeled based on corresponding cryo-EM density maps at 3.8-4.4Å resolution. Three protomers (a, b, c) are colored in red, blue and green, respectively. RBD locations are indicated. Particle percentage for each class in the data processing is also indicated, but it may not accurately reflect the conformation distribution of the S trimer in solution.

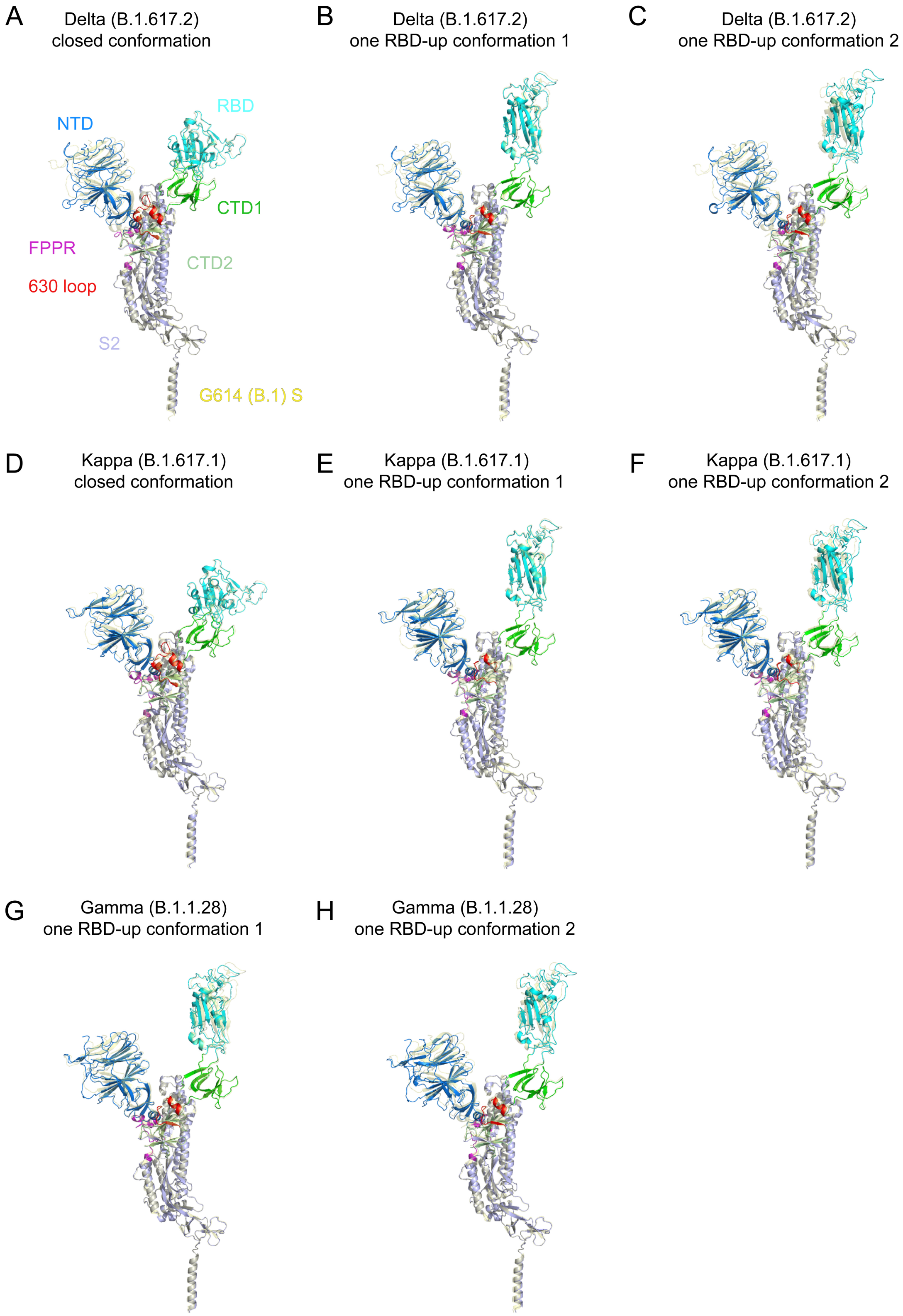
A
Delta (B.1.617.2)
closed conformation
NTD
RBD
CTD1
FPPR
CTD2
630 loop
S2
G614 (B.1) S
B
Delta (B.1.617.2)
one RBD-up conformation 1
C
Delta (B.1.617.2)
one RBD-up conformation 2
D
Kappa (B.1.617.1)
closed conformation
E
Kappa (B.1.617.1)
one RBD-up conformation 1
F
Kappa (B.1.617.1)
one RBD-up conformation 2
G
Gamma (B.1.1.28)
one RBD-up conformation 1
H
Gamma (B.1.1.28)
one RBD-up conformation 2

**Figure S15. Superposition of the variant trimer structures and the G614 structure.** (**A**)-(**C**) Side views of superposition of the closed conformation and two distinct one-RBD-up conformations of the Delta S (various colors) in ribbon diagram, aligned by the S2 portion with the closed prefusion structure and the one-RBD-up conformation of the G614 S (yellow), respectively. The positions of the RBD-down and three different RBD-up conformations are indicated. (**D**)-(**F**) Comparison of the Kappa S (various colors) and G614 trimer (yellow) structures in the corresponding conformations, when aligned by S2. (**G**) and (**H**) Comparison of the Gamma S (various colors) and G614 trimer (yellow) structures in the one-RBD-up conformation, when aligned by S2.

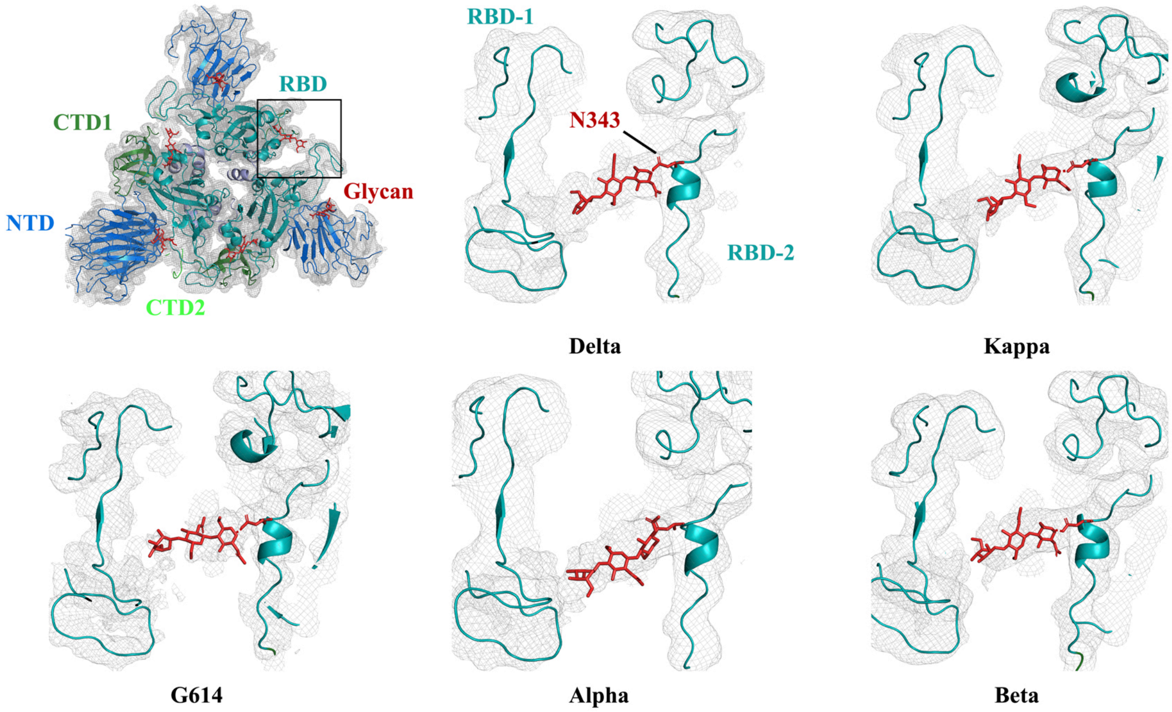


**Figure S16. Comparison of the density of the N-linked glycan at Asn343 in the variant trimers.** The EM maps of G614, Delta, Kappa, Alpha and Beta trimers in the closed prefusion conformation are compared at the same resolution (3.6Å) and the same contour level.

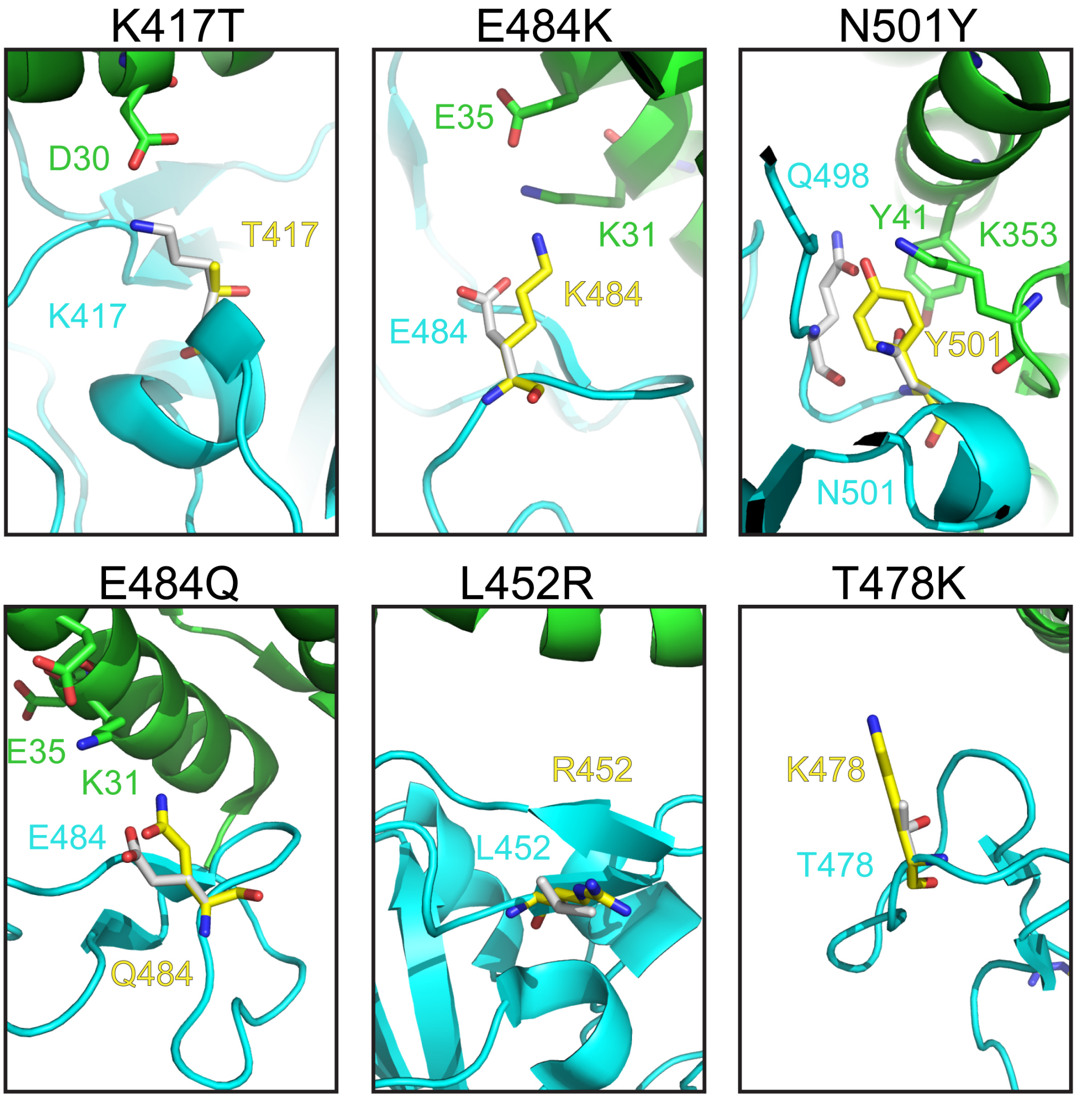


**Figure S17. Modeled interface between the RBD and ACE2.** The interface between ACE2 in ribbon diagram in green and RBD in cyan from the complex structure (PDB ID: 6M0J; ref(*36*)). Modeled K417T, E484K, N501Y, E484Q, L452R and T478K are shown as sticks. Panels E484K and N501Y were published previously in ref(*26*).

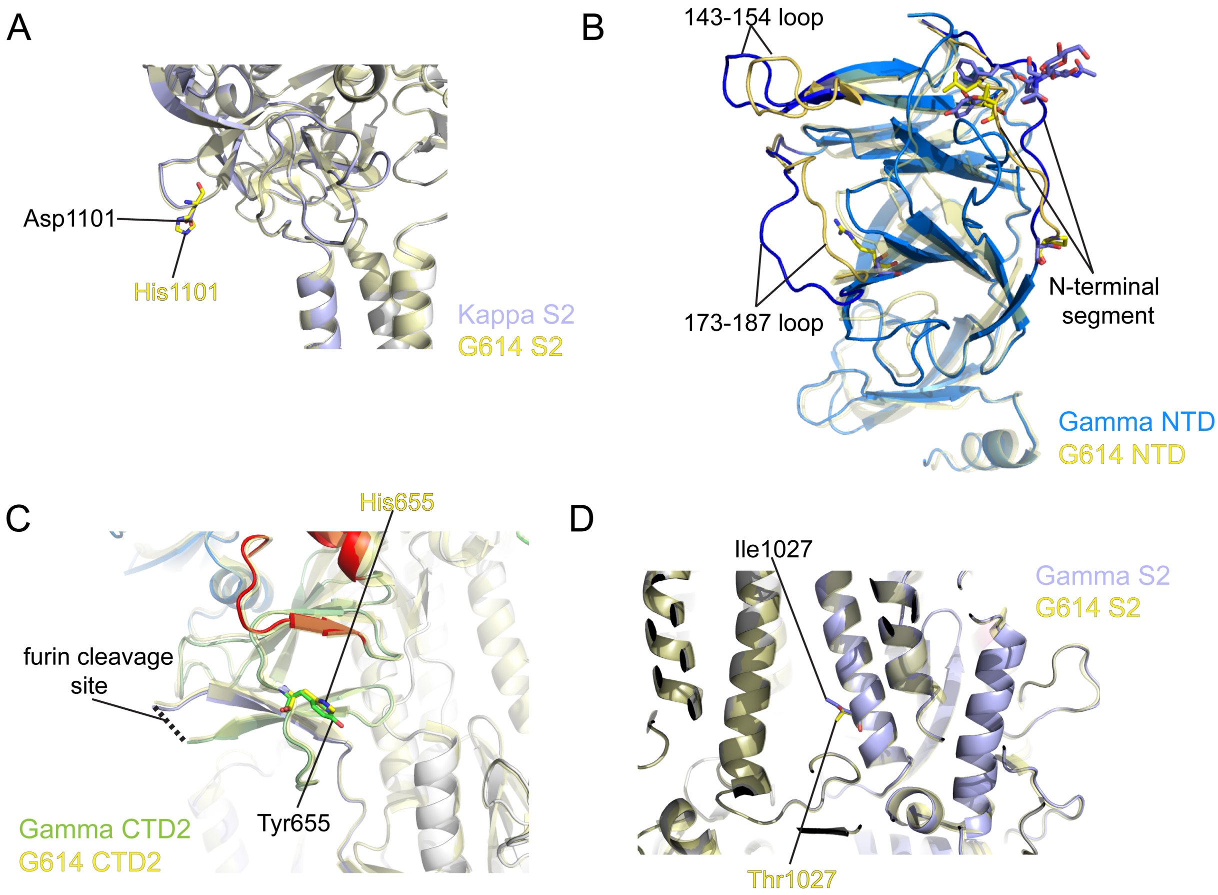


**Figure S18. Structural impact of the mutations in the variants**. (**A**) Superposition of the structure of the Kappa S trimer in ribbon representation with the structure of the G614 S in yellow, showing the regions near the mutation H1101D. (**B**) Superposition of the NTD structures of the Gamma (blue) and G614 (yellow) S trimers in the one-RBD-up conformation. Replacements of the N-terminal segment, 143-154 and 173-187 loops are indicated. (**C**) and (**D**) Superposition of the structure of the Gamma S trimer in ribbon representation with the structure of the G614 S in yellow aligned by S2, showing the region near mutations H655Y and T1027I. All mutations are shown as sticks.

### Table S1. Binding constants of S-ACE2 interaction

| | | $K_D$ (M) | $K_D$ Error | ka (1/Ms) | ka2 | ka Error | ka2 Error | kdis (1/s) | kdis2 | kdis Error | kdis2 Error |
|---|---|---|---|---|---|---|---|---|---|---|---|
| **ACE2-Fc (RBD)** | G614 S | 1.96E-08 | 2.52E-09 | 2.64E+04 | 6.00E-01 | 2.29E+03 | 7.68E+00 | 5.18E-04 | 8.98E-02 | 4.91E-05 | 1.13E+00 |
| | Gamma S | 2.41E-09 | 2.76E-10 | 9.45E+04 | 2.09E+01 | 4.95E+03 | 2.16E+00 | 2.28E-04 | 7.22E-01 | 2.31E-05 | 1.70E-01 |
| | Kappa S | 7.51E-09 | 8.99E-10 | 5.14E+04 | 4.29E-01 | 3.36E+03 | 5.44E-01 | 3.86E-04 | 3.85E-02 | 3.88E-05 | 5.59E-02 |
| | Delta S | 1.76E-07 | 5.64E-08 | 2.62E+04 | 1.06E-01 | 1.13E+03 | 4.38E-02 | 4.60E-03 | 1.83E-03 | 1.46E-03 | 4.41E-04 |
| | G614 RBD | 1.55E-08 | 1.30E-09 | 2.83E+04 | 2.27E-03 | 2.14E+03 | 1.07E+03 | 4.40E-04 | 2.77E+00 | 1.60E-05 | 1.30E+06 |
| | Gamma RBD | 8.76E-09 | 5.67E-10 | 3.95E+04 | 4.08E+00 | 1.21E+03 | 8.31E+00 | 3.46E-04 | 5.48E-01 | 1.97E-05 | 1.11E+00 |
| | Kappa RBD | 7.34E-09 | 2.82E-10 | 4.02E+04 | 3.31E+00 | 1.10E+03 | 1.05E+02 | 2.95E-04 | 1.07E+00 | 7.95E-06 | 3.38E+01 |
| | Delta RBD | 7.66E-09 | 5.93E-10 | 2.64E+04 | 4.17E-02 | 1.72E+03 | 9.46E-01 | 2.02E-04 | 9.36E-02 | 8.40E-06 | 2.10E+00 |
| **Monomeric ACE2 (RBD)** | G614 S | 2.88E-07 | 5.13E-09 | 1.00E+05 | | 1.71E+03 | | 2.89E-02 | | 1.48E-04 | |
| | Gamma S | 1.48E-08 | 1.92E-10 | 2.20E+05 | | 2.75E+03 | | 3.26E-03 | | 1.10E-05 | |
| | Kappa S | 1.60E-07 | 4.00E-09 | 9.71E+04 | | 2.31E+03 | | 1.55E-02 | | 1.22E-04 | |
| | Delta S | 2.08E-07 | 6.04E-09 | 7.38E+04 | | 2.05E+03 | | 1.53E-02 | | 1.33E-04 | |
| | G614 RBD | 2.56E-07 | 6.44E-09 | 8.15E+04 | | 1.96E+03 | | 2.09E-02 | | 1.55E-04 | |
| | Gamma RBD | 1.24E-07 | 1.96E-09 | 5.68E+04 | | 8.58E+02 | | 7.06E-03 | | 3.16E-05 | |
| | Kappa RBD | 7.63E-08 | 1.38E-09 | 1.59E+05 | | 2.75E+03 | | 1.21E-02 | | 6.42E-05 | |
| | Delta RBD | 1.99E-07 | 4.46E-09 | 6.16E+04 | | 1.32E+03 | | 1.23E-02 | | 8.06E-05 | |
| **C63C8 (RBD-1)** | G614 S | 7.05E-09 | 4.38E-11 | 8.92E+04 | | 3.64E+02 | | 6.29E-04 | | 2.95E-06 | |
| | Gamma S | 2.78E-09 | 1.56E-11 | 2.03E+05 | | 8.41E+02 | | 5.65E-04 | | 2.13E-06 | |
| | Kappa S | 1.90E-10 | 1.12E-11 | 1.31E+05 | | 5.07E+02 | | 2.50E-05 | | 1.47E-06 | |
| | Delta S | 4.76E-09 | 1.97E-11 | 1.95E+05 | | 6.97E+02 | | 9.29E-04 | | 1.94E-06 | |
| **G32B6 (RBD-2)** | G614 S | 7.05E-10 | 2.13E-11 | 2.12E+05 | 2.76E+00 | 2.73E+03 | 1.09E-01 | 1.49E-04 | 3.22E-01 | 4.08E-06 | 2.31E-02 |
| | Gamma S | N.D. | N.D. | N.D. | N.D. | N.D. | N.D. | N.D. | N.D. | N.D. | N.D. |
| | Kappa S | *3.43E-09*[a] | *2.09E-09* | | | | | | | | |
| | Delta S | 1.71E-09 | 2.10E-11 | 2.44E+05 | 3.22E+00 | 1.58E+03 | 5.29E+00 | 4.19E-04 | 6.43E-01 | 4.35E-06 | 1.05E+00 |
| **C12A2 (RBD-2)** | G614 S | 9.00E-10 | 1.69E-11 | 2.01E+05 | 1.52E-01 | 1.94E+03 | 1.42E-02 | 1.81E-04 | 2.27E-02 | 2.93E-06 | 2.18E-03 |
| | Gamma S | N.D. | N.D. | N.D. | N.D. | N.D. | N.D. | N.D. | N.D. | N.D. | N.D. |
| | Kappa S | *1.12E-08* | *1.33E-09* | | | | | | | | |
| | Delta S | 1.14E-09 | 1.65E-11 | 1.69E+05 | 1.10E-01 | 1.43E+03 | 1.39E-02 | 1.92E-04 | 2.73E-02 | 2.26E-06 | 3.61E-03 |
| **C63C7 (RBD-3)** | G614 S | 4.64E-09 | 1.97E-11 | 9.09E+04 | | 3.04E+02 | | 4.22E-04 | | 1.10E-06 | |
| | Gamma S | 2.86E-09 | 1.08E-11 | 1.65E+05 | | 4.44E+02 | | 4.72E-04 | | 1.25E-06 | |
| | Kappa S | 1.18E-10 | 9.51E-12 | 1.04E+05 | | 3.06E+02 | | 1.23E-05 | | 9.87E-07 | |
| | Delta S | 2.36E-09 | 7.20E-12 | 1.94E+05 | | 4.01E+02 | | 4.58E-04 | | 1.03E-06 | |
| **C63D3 (RBD-3)** | G614 S | 6.87E-09 | 4.45E-11 | 2.02E+05 | | 1.19E+03 | | 1.39E-03 | | 3.75E-06 | |
| | Gamma S | 6.29E-09 | 3.40E-11 | 2.81E+05 | | 1.41E+03 | | 1.77E-03 | | 3.61E-06 | |
| | Kappa S | *1.84E-08* | *1.07E-08* | | | | | | | | |
| | Delta S | 6.82E-09 | 3.79E-11 | 1.63E+05 | | 8.10E+02 | | 1.11E-03 | | 2.80E-06 | |
| **C12C9 (NTD-1)** | G614 S | 3.50E-09 | 1.91E-11 | 7.88E+04 | | 2.94E+02 | | 2.75E-04 | | 1.10E-06 | |
| | Gamma S | *1.10E-08* | *6.48E-09* | | | | | | | | |
| | Kappa S | *3.31E-09* | *6.32E-10* | | | | | | | | |
| | Delta S | N.D. | N.D. | N.D. | | N.D. | | N.D. | | N.D. | |
| **C83B6 (NTD-1)** | G614 S | 1.07E-09 | 5.27E-12 | 1.86E+05 | | 3.60E+02 | | 1.99E-04 | | 9.03E-07 | |
| | Gamma S | N.D. | N.D. | N.D. | | N.D. | | N.D. | | N.D. | |
| | Kappa S | *~1.13E-07* | *N.A.* | | | | | | | | |
| | Delta S | N.D. | N.D. | N.D. | | N.D. | | N.D. | | N.D. | |
| **C81D6 (NTD-2)** | G614 S | 3.99E-09 | 4.74E-11 | 7.25E+04 | | 6.25E+02 | | 2.89E-04 | | 2.36E-06 | |
| | Gamma S | 1.91E-09 | 5.92E-11 | 4.84E+04 | | 6.81E+02 | | 9.25E-05 | | 2.56E-06 | |
| | Kappa S | *4.16E-09* | *1.49E-09* | | | | | | | | |
| | Delta S | 6.87E-10 | 2.25E-11 | 7.02E+04 | | 4.30E+02 | | 4.83E-05 | | 1.56E-06 | |
| **C163E6 (S2-2)** | G614 S | *6.47E-09* | *2.38E-09* | | | | | | | | |
| | Gamma S | *2.04E-08* | *1.37E-08* | | | | | | | | |
| | Kappa S | *3.49E-08* | *5.11E-09* | | | | | | | | |
| | Delta S | *1.03E-08* | *1.75E-09* | | | | | | | | |

[a]Values in italic are derived from steady-state fitting.

**Table S2. Neutralization of the SARS-CoV-2 variants**

| Antibody/ACE2 construct | Neutralization titer (µg/ml) | | | | | | | |
|---|---|---|---|---|---|---|---|---|
| | B.1 (G614) | | Gamma | | Kappa | | Delta | |
| | $IC_{50}$ | $IC_{80}$ | $IC_{50}$ | $IC_{80}$ | $IC_{50}$ | $IC_{80}$ | $IC_{50}$ | $IC_{80}$ |
| C63C8 (RBD-1) | 8.030 | 27.794 | 4.378 | 24.127 | 5.619 | 25.555 | 4.563 | 19.939 |
| C12A2 (RBD-2) | 0.014 | 0.071 | >50 | >50 | >50 | >50 | 0.063 | 0.339 |
| G32B6 (RBD-2) | 0.018 | 0.051 | >50 | >50 | >50 | >50 | 0.015 | 0.048 |
| C63C7 (RBD-3) | 4.729 | 29.985 | 3.273 | 18.781 | 4.532 | 20.257 | 7.704 | 34.343 |
| C63D3 (RBD-3) | 37.186 | >50 | 30.614 | >50 | >50 | >50 | 46.126 | >50 |
| C12C9 (NTD-1) | 0.019 | 43.34 | 0.341 | >50 | 22.939 | >50 | >50 | >50 |
| C83B6 (NTD-1) | 0.105 | >50 | >50 | >50 | >50 | >50 | >50 | >50 |
| C81D6 (NTD-2) | >50 | >50 | >50 | >50 | >50 | >50 | >50 | >50 |
| C163E6 (S2-2) | >50 | >50 | >50 | >50 | >50 | >50 | >50 | >50 |
| ACE2-T27W-Fd | 0.079 | 0.568 | 0.036 | 0.180 | 0.045 | 0.147 | 0.022 | 0.054 |
| Positive serum pool_2 (1/x) | 1,922 | 426 | 58 | 23 | 1,046 | 96 | 2,300 | 150 |
| Normal Human Serum (1/x) | <20 | <20 | <20 | <20 | <20 | <20 | <20 | <20 |

### Table S3. Cryo-EM statistics.

**EM data collection and reconstruction statistics**

| Protein | Full-length S of Delta variant | | | Full-length S of Gamma variant | | Full-length S of Kappa variant | | |
|---|---|---|---|---|---|---|---|---|
| Microscope | Titan Krios | | | Titan Krios | | Titan Krios | | |
| Voltage(kV) | 300 | | | 300 | | 300 | | |
| Detector | Gatan K3 | | | Gatan K3 | | Gatan K3 | | |
| Magnification(nominal) | 105,000 | | | 105,000 | | 105,000 | | |
| Energy filter slit width (eV) | 20 | | | 20 | | 20 | | |
| Calibrated pixel size (Å/pix) | 0.825 | | | 0.825 | | 0.825 | | |
| Exposure rate (e$^-$/pix/sec) | 20.24 | | | 20.69/20.63/27.13 | | 21.12/20,101 | | |
| Frames per exposure | 50 | | | 51/51/50 | | 51/51 | | |
| Total electron exposure (e$^-$/Å$^2$) | 51.48 | | | 54.72/54.56/53.4 | | 51.63/51.151 | | |
| Exposure per frame (e$^-$/Å$^2$) | 1.03 | | | 1.073/1.07/1.06 | | 1.012/1.003 | | |
| Defocus range (μm) | -0.8,-2.2 | | | -0.8, -2.3 | | -0.8, -2.2 | | |
| Automation software | SerialEM | | | SerialEM | | SerialEM | | |
| # of Micrographs used | 20,274 | | | 25,424/32,569/29,163 | | 22,019/17,314 | | |
| Particles extracted | 1,830,328 | | | 1,652,420/2,757,190/1,893,863 | | 1,199,999/1,718,600 | | |
| Particles after 2D classification | 1,386,630 | | | 1,564,938/1,904,078/1,788,000 | | 1,112,384/1,649,296 | | |
| Class | Closed | RBD-up 1 | RBD-up 2 | RBD-up 1 | RBD-up 2 | Closed | RBD-up 1 | RBD-up 2 |
| Total # of refined particles | 94,680 | 191,067 | 25,370 | 69,302 | 36,346 | 123,193 | 81,717 | 21,830 |
| Symmetry imposed | C3 | C1 | C1 | C1 | C1 | C3 | C1 | C1 |
| Estimated accuracy of translations/rotations | 0.87/1.96 | 1.23/2.45 | 2.35/4.24 | 1.73/3.11 | 2.27/3.80 | 1.04/2.29 | 1.66/3.24 | 1.86/3.36 |
| Map sharpening B-factor | -93.6 | -111.9 | -92.1 | -124.6 | -153.7 | -103.6 | -121.8 | -130.3 |
| Unmasked Resolution at 0.5/0.143 FSC (Å) | 3.9/3.4 | 4.4/3.8 | 8.9/7.3 | 7.1/4.3 | 8.4/6.1 | 4.0/3.6 | 6.4/4.0 | 8.5/5.9 |
| Masked resolution at 0.5/0.143 FSC (Å) | 3.6/3.1 | 3.9/3.4 | 7.3/4.3 | 4.5/3.8 | 6.9/4.4 | 3.6/3.1 | 4.2/3.7 | 7.4/4.3 |

**Model refinement and validation statistics**

| Class | Closed | RBD-up 1 | RBD-up 2 | RBD-up 1 | RBD-up 2 | Closed | RBD-up 1 | RBD-up 2 |
|---|---|---|---|---|---|---|---|---|
| PDB | | | | | | | | |
| Composition | | | | | | | | |
| Amino acids | 3315 | 3287 | 3269 | 3230 | 3300 | 3330 | 3270 | 3270 |
| Glycans | 57 | 57 | 57 | 60 | 60 | 57 | 57 | 57 |
| RMSD bonds (Å) | 0.016 | 0.014 | 0.015 | 0.013 | 0.013 | 0.013 | 0.014 | 0.015 |
| RMSD angles (◦) | 2.11 | 1.85 | 1.92 | 1.81 | 1.83 | 1.80 | 1.78 | 1.88 |
| Mean B-factors | | | | | | | | |
| Amino acids | 62 | 64 | 64 | 62 | 62 | 62 | 64 | 64 |
| Glycans | 92 | 88 | 88 | 88 | 88 | 92 | 88 | 87 |
| Ramachandran | | | | | | | | |
| Favored (%) | 92.33 | 92.63 | 91.68 | 92.16 | 91.96 | 93.09 | 93.07 | 91.58 |
| Allowed(%) | 6.48 | 6.64 | 7.42 | 7.17 | 7.16 | 6.06 | 6.40 | 7.77 |
| Outliers(%) | 1.19 | 0.74 | 0.90 | 0.67 | 0.89 | 0.85 | 0.53 | 0.65 |
| Rotamer outliers (%) | 5.92 | 4.40 | 3.86 | 2.07 | 1.84 | 2.41 | 1.89 | 2.38 |
| Clash score | 3.70 | 2.39 | 3.42 | 1.85 | 3.43 | 1.63 | 2.75 | 2.67 |
| C-beta outliers (%) | 1.23 | 0.81 | 1.01 | 048 | 0.49 | 0.35 | 0.33 | 0.39 |
| CaBLAM outliers (%) | 2.86 | 3.01 | 3.57 | 2.75 | 2.99 | 2.57 | 2.60 | 3.07 |
| CC (mask) | 0.78 | 0.76 | 0.61 | 0.69 | 0.63 | 0.79 | 0.73 | 0.63 |
| CC (volume) | 0.78 | 0.76 | 0.60 | 0.69 | 0.63 | 0.78 | 0.73 | 0.62 |
| MolProbity score | 2.22 | 1.97 | 2.08 | 1.67 | 1.82 | 1.65 | 1.72 | 1.84 |
| EMRinger score | 3.65 | 2.61 | 0.88 | 1.76 | 0.74 | 3.00 | 2.27 | 0.73 |
